# Organism-scale annotation with Pan-human Azimuth

**DOI:** 10.64898/2026.07.16.738997

**Authors:** Sourav Sarkar, Zhuoyan Li, Gesmira Molla, Anagha Shenoy, Brian Zhang, David Collins, Nicole Vasilevsky, Joseph P. Gaut, Aleix Puig-Barbe, Andreas Bueckle, David Osumi-Sutherland, Katy Börner, Sanjay Jain, Rahul Satija

## Abstract

Single-cell atlases now span many human tissues, but inconsistent annotations across studies limit their utility as a unified reference. We introduce Pan-human Azimuth, a supervised neural network that maps human cells from diverse tissues and datasets onto a single hierarchical organism-scale typology. Developed through NIH HuBMAP, the model is trained on a uniquely curated corpus designed to maximize diversity across tissues and technologies while enforcing uniform, interpretable annotations and stringent quality control. The use of a single organism-wide reference enables us to map tens of millions of cells in the Tabula Sapiens and scBaseCamp repositories, perform cross-tissue comparisons across thousands of samples, and identify striking tissue specialization among fibroblast states. Pan-human Azimuth naturally extends to annotating spatial transcriptomic data, recovering canonical kidney cortical structures and distinguishing glomerular states consistent with expert pathology. We release Pan-human Azimuth alongside cloud, R, and Python interfaces to facilitate standardized organism-wide single-cell analysis.

## 1 INTRODUCTION

The emergence of single-cell RNA sequencing (scRNA-seq) ^1–4^ has enabled the high-resolution discovery of cell types and the characterization of cellular behavior across diverse biological contexts. This has led to substantial community-driven efforts, for example via the Human Cell Atlas (HCA) ^5^ and the NIH Human Biomolecular Atlas Program (HuBMAP) ^6,7^, to construct comprehensive molecular atlases of human organs and tissues. As these reference maps reach maturity, they enable a transition in analytical workflows from the unsupervised analysis of isolated datasets toward reference-based supervised analysis that leverages the cumulative knowledge present in cellular atlases. Supervised reference-mapping approaches not only boost statistical power to identify rare and subtle cell states, but also facilitate reproducible and uniform workflows across disparate samples and studies. Despite this promising vision, however, implementing reference-based analysis on an organismal scale remains a formidable challenge.

Previous strategies for reference-based cell annotation have largely taken two distinct forms, each operating at a different biological scale. The first involves building individual organ-specific references trained to predict cell types within a single tissue^8–13^. While effective for harmonizing datasets from the same organ, these references do not enforce consistent annotations across tissues and are not designed to reveal biological patterns that emerge only within a unified, organism-wide framework. Single-cell foundation models ^14–18^ offer an alternative by using self-supervised learning to embed cells in a low-dimensional space, enabling annotation through nearest-neighbor matching. However, these models generally rely on author-provided annotations aggregated from thousands of studies, where inconsistent naming conventions create “label fragmentation” ^19–21^, which limits the accuracy and interpretability of downstream results. While foundation models often emphasize the size and scale of training data, recent work has highlighted how diversity, quality, and harmonization of training data are key determinants of model performance ^22^.

To address these challenges, we developed Pan-human Azimuth as part of the NIH HuBMAP project, establishing a scalable framework for unified organism-wide cell annotation under a harmonized hierarchical typology. We assembled a training corpus designed to capture broad biological and technical diversity across tissues, studies, donors, and profiling platforms. Central to this effort was a uniform curation process that placed every cell within a consistent hierarchical cell type tree. By combining this standardized taxonomy with stringent error checking and quality control, we prioritized the accuracy and robustness of the labels used for model training. This curated resource enabled us to train a supervised neural network-based hierarchical classifier optimized for multi-resolution cell annotation of new datasets. For each cell, Pan-human Azimuth returns both an interpretable cell type identity and a calibrated confidence score, allowing users to assess predictions across different levels of annotation granularity.

We demonstrate how the key features of our framework enable organism-scale analysis of human single-cell data. Joint training allows information from one tissue to improve annotation in another, including for rare or sparsely represented populations; for example, high-resolution T-cell states learned from immune-rich tissues support more refined immune annotation in the lung. The shared hierarchy also allows entire organism-wide atlases to be mapped consistently in a single run, including Tabula Sapiens ^23,24^ and 85.9 million human cells from scBaseCamp ^25^. This enables robust cross-tissue analyses across thousands of samples that cannot be performed when using tissue-specific references with different annotation schemes and levels of resolution. Using this framework, we show that fibroblast state composition reproducibly predicts tissue of origin across thousands of samples, revealing tissue-specialized patterns that would otherwise be obscured by study-specific nomenclature. Finally, Pan-human Azimuth extends naturally to high-resolution spatial transcriptomic data: when applied to Visium HD kidney samples ^26,27^, it recovers canonical cortical organization and supports classification of healthy and sclerosed glomeruli in agreement with expert pathologist annotations. Together, these results establish Panhuman Azimuth as a powerful and interpretable resource for organism-wide annotation and comparative analysis of human single-cell data.

## RESULTS

Here we introduce Pan-human Azimuth, an organism-wide annotation resource for human single-cell and single-nucleus RNA-seq data. To construct Pan-human Azimuth, we assembled a harmonized training corpus spanning 23 tissues and more than 27 million cells from five major resources. We systematically re-annotated these data within a single hierarchical framework, converting fragmented and heterogeneous cell identity labels into consistent organism-wide annotations. We then used this resource to train a deep-learning-based hierarchical classifier that maps gene expression count matrices onto a structured cell type typology. The model returns annotations at eight levels of resolution, from broad lineages to fine transcriptional subtypes, together with calibrated confidence scores. Below, we describe the construction of Pan-human Azimuth, benchmark its performance against existing organism-scale annotation strategies, and demonstrate its ability to uniformly annotate human cells across tissues, technologies, and biological contexts.

### Construction of the Training Dataset

To build a robust foundation for a generalizable human cell type classification model, we assembled a large and diverse training corpus spanning 23 tissues and more than 27 million cells (Fig.1A). This corpus integrated six Azimuth^9^ references (e.g., heart, liver, spleen) ^28–46^, 21 DISCO references (including adrenal gland, kidney, lung) ^11^, 123 datasets from CELLxGENE ^47,48^, five tissues from GTEx ^49,50^, and 215 datasets from HuBMAP ^6,7,51^. As published single-cell datasets may potentially be represented in multiple public repositories, we developed a random projection-based approach to identify and remove duplicated submissions (Supplementary Methods). We then performed systematic hierarchical re-annotation of all cells, as described further below, to harmonize source annotations, enforce ontological consistency across tissues and datasets, and improve the accuracy of labels used for supervised model training (Fig.1B).

**Figure 1.**
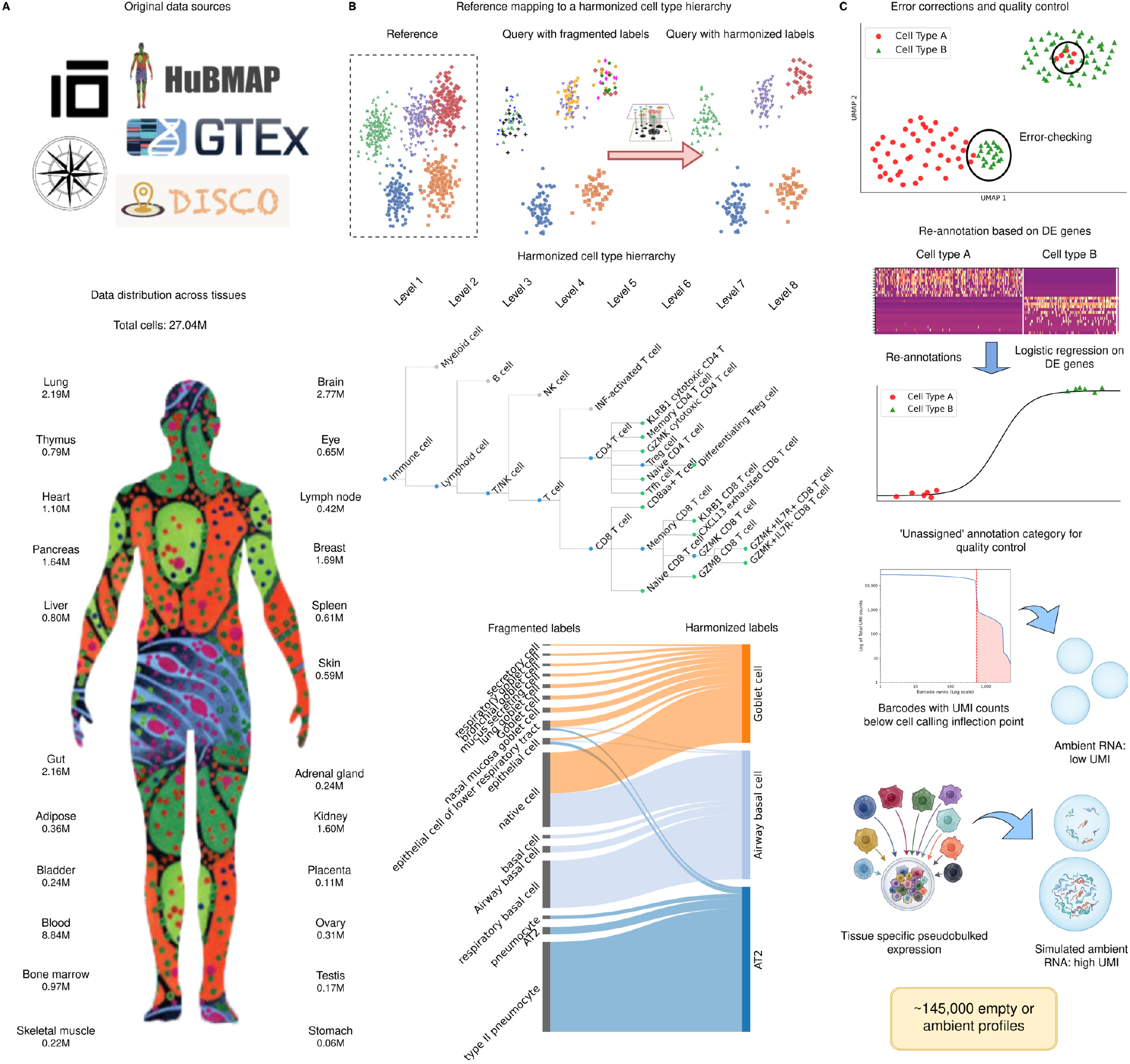
Preparation of the Pan-human Azimuth training dataset. (A) Collection of whole-body human scRNA-seq data: (Top) Data for the training corpus was collected from CZI CELLxGENE, HuBMAP, Azimuth, GTEx, and DISCO. (Bottom) A total of approximately 27.04 million cellular profiles from 23 tissues across the body were collected. Schematic of the human body is adapted from the NIH HuBMAP logo. (B) Harmonization of the cell type hierarchy through reference-based label transfer: (Top) Cell types were projected onto a common hierarchical tree using label transfer from tissue-specific references. (Middle) The hierarchical cell type tree provides annotations for each cell at varying levels of resolution (Bottom) A direct transfer of author-provided labels from disparate references leads to fragmentation of labels (as seen on the left-hand-side of the Sankey diagram). The projection onto a common hierarchical tree harmonizes the labels. (C) Annotation refinement and quality control. Top: Following initial label transfer, dataset-specific high-resolution clusters were used to identify candidate annotation inconsistencies for error checking. Middle: Candidate cells were re-evaluated using logistic regression models, and cells were reassigned when their expression profiles more supported an alternative identity. Bottom: Low-UMI droplets and simulated ambient-RNA profiles generated from tissue-specific pseudobulks were included as approximately 145,000 negative training examples labeled “Unassigned,” enabling the model to identify low-quality or non-cellular profiles during inference.

To anchor all annotations in a common framework, we used a single organism-wide cell type tree. We built on the DISCO cell type tree, which is organism-wide in scope and data-driven, having been derived directly from scRNA-seq data rather than imposed from prior anatomical classifications ^11^. As part of the NIH Human Biomolecular Atlas Program (HuBMAP), we made two refinements. First, we introduced targeted modifications to the tree structure to incorporate cell types and distinctions that have emerged from recent large-scale atlas efforts. Second, we systematically built a one-to-one mapping from each node and leaf in the resulting tree to the Cell Ontology ^21^, a widely adopted controlled vocabulary for cell type annotation. The resulting tree provides a unified interpretive framework for all data in our corpus while also enabling users to translate to established ontology standards used across the field.

Systematic re-annotation of the training data proceeded in three steps (Fig.1B-C). First, for each of the 23 tissues, we selected an appropriate reference, typically DISCO or Azimuth, but in some cases a high-quality standalone atlas such as the Human Brain Cell Atlas^52^. We manually assigned each cell type label in these references onto the refined cell type tree, resolving synonymous and overlapping labels that accumulate across independent studies and generating a clean, non-redundant label space shared across tissues. We then used our previously developed anchor-based supervised label-transfer framework ^53^ to annotate all ~27 million query cells.

Although resources such as CELLxGENE provide Cell Ontology mappings for many datasets, the resolution and granularity of these annotations vary widely across studies, reflecting differences in expertise, biological focus, and labeling conventions among contributing groups. For example, CELLxGENE respiratory epithelial annotations include multiple labels such as “respiratory goblet cell”, “mucus secreting cell”, “mucus secreting goblet cell”, “secretory cell”, and “native cell”, which are harmonized in our framework to “Goblet cell” in all cases (Fig.1B bottom). By deriving annotations through reference mapping onto a single shared tree, we ensured that query datasets were labeled uniformly and at consistent resolution, independent of the terminology used in the original studies.

Second, we implemented a local annotation refinement step to improve annotation accuracy after initial label transfer (Supplementary Methods). We independently clustered each query dataset at high resolution and compared each cell’s assigned label with the majority label of its local cluster, flagging discrepancies as potential annotation errors (Fig.1C top; Supplementary Methods). For each flagged case, we then trained a pairwise logistic-regression classifier using genes that distinguished the assigned label from the cluster-majority label (Fig.1C middle). This classifier provided an expression-based test of whether the cell’s transcriptomic profile better supported its original annotation or the alternative local annotation. Cells with stronger support for the alternative label were reassigned as a form of error-correction. Applied across the full training corpus, this refinement reassigned approximately 7% of cells, resolving local inconsistencies and improving the reliability of labels used for supervised model training.

Third, we implemented an independent annotation quality-control step to retain only high-confidence cells for model training (Supplementary Methods). For each cell type assignment, we estimated annotation reliability using one-versus-all logistic-regression models trained on 100 positive and negative reference-derived marker genes for each cell type. The resulting probabilities provided continuous confidence scores for the assigned identity of each cell within the training corpus. We removed low-confidence cells with scores below 0.75, then selected approximately 9.65 million cells representing a balanced mixture of cell types and tissues for inclusion in the final training dataset. Together, these three steps represent a scalable approach that applies the principles of manual QC, error-checking, and harmonized labeling to create an organism-scale training corpus.

Lastly, we reasoned that, in addition to accurately annotated cells, labeled empty droplets and ambient RNA profiles could provide negative examples that enable built-in quality control by training the model to recognize low-quality data as its own category to be filtered out (Fig.1C bottom). To collect empty droplets, we downloaded publicly available 10x Genomics datasets and extracted barcodes with UMI counts below the inflection point used for cell calling. We also simulated ambient RNA profiles from negative binomial distributions derived from tissue-specific pseudobulk expression profiles, reasoning that these examples would complement empty droplets by capturing higher-UMI artifacts that nevertheless do not represent true cells. In total, we generated approximately 145,000 empty-droplet and ambient-RNA profiles, labeled them as “Unassigned,” and added them to the training dataset. Taken together, these steps produced a training dataset that spans a broad range of tissues, studies, donors, and technologies, with harmonized annotations, a unified ontological structure, and multiple layers of quality control. These features make it well-suited for training a broadly generalizable human cell classification model.

### Pan-human Azimuth Neural Network Architecture and Training

We next used this curated training dataset to train a deep-learning classifier for annotating cells in new datasets. We constructed a model consisting of three main components: an embedding module that converts sparse single-cell count data into a compact low-dimensional representation, a set of hierarchically organized classification modules, and a layer of confidence calibration models that align predicted probabilities at each hierarchical level with empirical accuracy (Fig.2A,C,E). The network first passes each input expression profile through a series of fully connected layers, compressing it into a 128-dimensional embedding that captures transcriptional features most informative for distinguishing cellular annotations. This compact representation serves as a shared input to all downstream classification modules. This space also serves as a unified embedding coordinate system across datasets, and can be used as input to generate two-dimensional visualizations ^54^.

**Figure 2.**
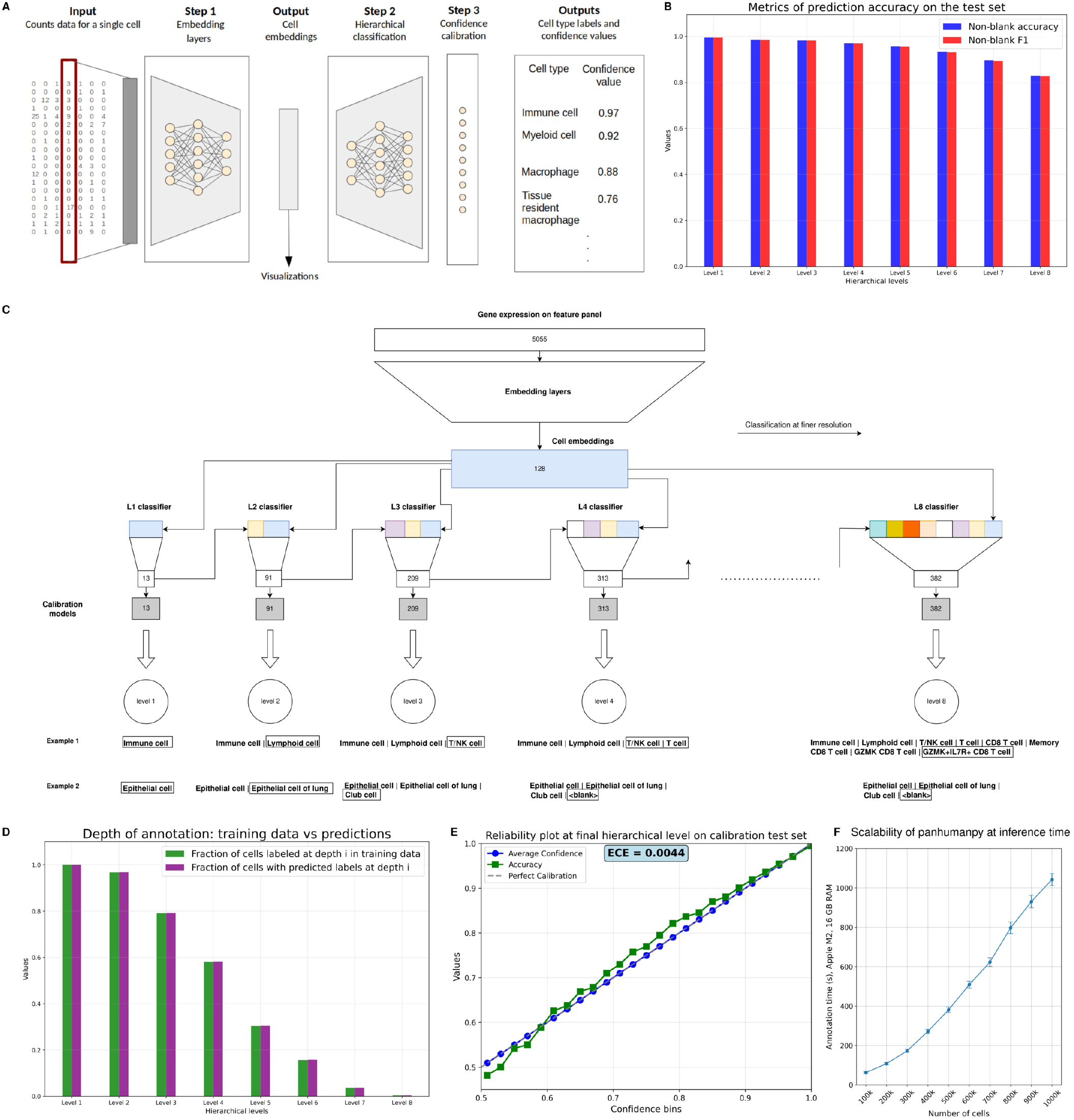
Overview of the Pan-human Azimuth Neural Network. (A) Overview of the Pan-human Azimuth architecture. Gene-expression profiles from an scRNA-seq dataset are first compressed through a series of embedding layers into a low-dimensional representation. A hierarchical classifier then assigns each cell annotations at progressively finer levels of resolution, and post hoc calibration models provide confidence scores for predictions at each hierarchical level. (B) Annotation performance across the hierarchy on a held-out test set. “Non-blank accuracy” denotes accuracy calculated only among cells with a non-blank ground-truth label at the corresponding level. “Non-blank F1” similarly denotes the F1 score after excluding cells whose annotation hierarchy terminates above that level. (C) Detailed architecture of the hierarchical classifier. Each cell’s 128-dimensional embedding is passed through a nested series of multilayer perceptrons, with one classification head responsible for each hierarchical level. The number of possible labels increases with depth, reflecting the greater complexity of finer-resolution annotation. (D) Comparison of the maximum annotation depth assigned to each cell in the ground-truth labels and model predictions. (E) Confidence calibration at the deepest available annotation level for each cell in the calibration test set. Following calibration, predicted confidence closely matches empirical accuracy. (F) Inference scalability of panhumanpy. Query datasets ranging from 100,000 to 1 million cells were annotated on a MacBook Air with an Apple M2 8-core processor and 16 GB of memory using a batch size of 8,192. Total inference time increased approximately linearly with dataset size, while throughput remained stable, demonstrating efficient scaling on modest hardware. Error bars represent the standard error across 10 runs.

The network next decodes these embeddings into a set of multi-level cell annotations using a set of hierarchically organized classification heads, each corresponding to a level in the cell type typology. For example, the first classifier (Level 1) predicts broad categories such as immune, epithelial, or stromal cells, while deeper classifiers (Levels 2–8) refine these categories into increasingly specialized subtypes (Fig.2B). Each classification head is a multilayer perceptron (MLP) that operates on the same 128-dimensional embedding, together with a compressed signal from the decoded annotation at the preceding levels so that fine-grained predictions are aware of the parent category and are thus encouraged to be hierarchically consistent. Each head also includes a “blank” output, allowing the model to abstain from deeper annotation when evidence for a more refined classification is insufficient, while still returning a confident label at an intermediate level. The use of separate classification heads allows the model to solve both broad and fine-grained annotation tasks simultaneously, despite varying degrees of difficulty. At each level we calculated the focal loss ^55^, a variant of cross-entropy that mitigates class imbalance and emphasizes harder-to-classify examples, and the overall training objective function was a weighted sum of the per-level losses that balances learning across the hierarchy (Supplementary Methods).

Although the model outputs softmax probabilities that provide a natural measure of prediction confidence, neural networks can produce probabilities that are mis-calibrated relative to their empirical accuracy^56^. To address this, we applied entropy-informed temperature scaling ^56^ as a post-hoc calibration step (Supplementary Methods). We applied this separately to each of the eight hierarchical classification heads, training on a set held out from the main classifier training, to align predicted probabilities with observed empirical performance. The resulting models convert softmax probabilities to calibrated confidence scores for each hierarchical annotation in each cell (Fig.2E).

After training (Supplementary Methods), the model achieved high accuracy and F1 scores evaluated on all non-trivial labels across all hierarchical levels (Fig.2B), with performance remaining strong even for rare or deeply nested subclasses (Fig.2D, Supplementary Fig.1B). Post-hoc calibration aligned predicted confidence scores closely with empirical accuracies (Fig.2E, Supplementary Fig.1C). Finally, since each cell is annotated independently, total inference time scaled approximately linearly with dataset size (Fig.2F), while throughput approached 1,000 cells/second on a personal computer (MacBook Air; Apple M2 chip, 16 GB memory).

To assess how model performance scaled with the size of the training data, we downsampled the training corpus and evaluated annotation accuracy across hierarchical levels. Pan-human Azimuth benefited strongly from additional training data at small dataset sizes, but began to saturate as the corpus grew (Supplementary Fig.1A). Accuracy improved rapidly early on, with the largest gains for deeper, higher-resolution annotations, then increased more gradually beyond five million cells. This is consistent with recent studies that analyze the role of training dataset sizes ^22^, and suggests that future improvements are unlikely to come simply from adding more cells from already well-represented contexts. Instead, gains will likely depend on expanding training diversity, such as additional tissues, developmental time points, or rare cell states.

### Performance evaluation and benchmarking

We next evaluated model performance and generalization using datasets containing new donors that were not included in training. We applied Pan-human Azimuth to the Tabula Sapiens v2 atlas^24^, which contains 1.1 million single-cell and single-nucleus profiles generated using both droplet- and plate-based technologies across 24 donors and 28 tissues. Tabula Sapiens v2 includes data from 9 new donors contributing approximately 600,000 cells released after Pan-human Azimuth was trained. We ran the full atlas in a single inference step, enabling cells from all tissues and donors to be annotated, embedded, and visualized together within a unified coordinate system (Fig.3A-C). Pan-human Azimuth returned high-confidence annotations across the atlas (Fig.3D), with a median calibrated confidence score of 0.95 at the most fine-grained level assigned to each cell. Looking across multiple tissues we did not observe any systematic decrease in confidence scores for v2 donors compared to v1 donors (Supplementary Fig.2A), supporting the model’s ability to generalize to unseen samples.

**Figure 3.**
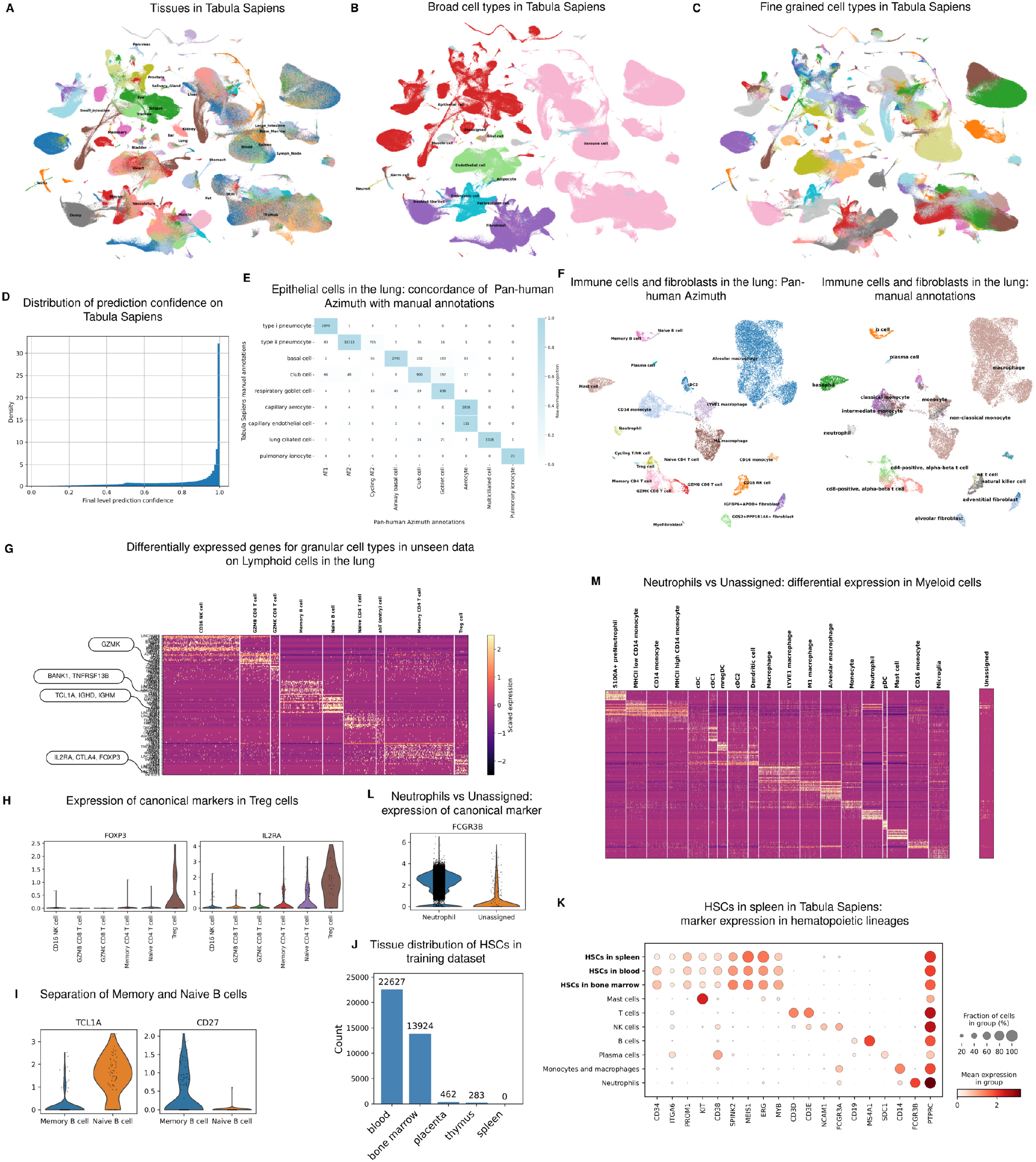
Annotating Tabula Sapiens with Pan-human Azimuth. (A) UMAP visualization of Pan-human Azimuth embeddings of Tabula Sapiens showing distribution across tissues. (B) UMAP visualization of the broad cell type annotations from Pan-human Azimuth. (C) UMAP visualization of finely resolved cell type annotations from Pan-human Azimuth. (D) Density distribution of prediction confidence values on the Tabula Sapiens dataset. (E) Confusion matrix showing the high degree of concordance between manual annotations from Tabula Sapiens and annotations from Pan-human Azimuth on epithelial cell types in the lung. (F) UMAP visualization of immune cells and fibroblasts in lung tissue data in Tabula Sapiens comparing Pan-human Azimuth-assigned annotations (left) with manual annotations (right). (G) Differential expression of markers in the set of Lymphoid cells in lung data from unseen donors in Tabula Sapiens v2 supports the accuracy of the finely resolved annotations by Pan-human Azimuth. (H) Expression of canonical markers *FOXP3* and *IL2RA* (*CD25*) in Treg cells in lung data from unseen donors. (I) Separation of Memory and Naive B cells in lung data from unseen donors based on expression of canonical markers *TCL1A* and *CD27*. (J) Distribution of HSCs across tissues in the Pan-human Azimuth training data. (K) HSCs identified in spleen samples from unseen Tabula Sapiens donors show expression of canonical hematopoietic stem and progenitor markers. (L) Evaluation of Pan-human Azimuth “Unassigned” calls among cells manually annotated as neutrophils. Cells classified as neutrophils by both approaches (left) express *FCGR3B*, whereas cells labeled “Unassigned” by Pan-human Azimuth (right) generally do not. (M) Expression of marker genes for the most abundant myeloid populations in Tabula Sapiens among cells labeled “Unassigned” by Pan-human Azimuth. The absence of coherent canonical marker expression supports the interpretation that these profiles are low quality or otherwise not confidently assignable to a single cell type.

We next compared Pan-human Azimuth annotations with the manually curated labels reported in the original Tabula Sapiens publication, focusing first on lung as a representative tissue (Supplementary Methods). For epithelial populations, manual and automated annotations were highly concordant (Fig.3E). This agreement is expected given the strong tissue specificity of many epithelial identities, and was further supported by differential gene expression analysis based on v2 donor annotations (Supplementary Fig.2B). In contrast, immune and stromal populations benefited from Pan-human Azimuth’s organism-wide training, which allowed the model to transfer shared immune-cell and fibroblast structure learned across tissues and assign more refined labels than the broad manual categories available within a single tissue. As a result, Pan-human Azimuth resolved substantial additional structure within lymphoid, myeloid, and stromal populations (Fig.3F-I). Canonical marker expression supported these higher-resolution assignments (Fig.3G), including enrichment of *GZMK* in effector CD8 T cells ^57^; *FOXP3, IL2RA*, and *CTLA4* in regulatory T cells ^58,59^ (Fig.3H); and *TCL1A* and *CD27* distinguishing naive and memory B cells ^60^ (Fig.3I). Pan-human Azimuth was also able to annotate HSC in the spleen of a Tabula Sapiens v2 donor. This was based entirely on patterns learned from other tissues such as blood and bone marrow (Fig.3J-K), as splenic HSC are known drivers of splenic hematopoiesis ^61^, but are very rare and were entirely missing from our training data. Together, these results show that Pan-human Azimuth recapitulates and extends expert annotations for tissue-specific and shared populations.

We next assessed whether Pan-human Azimuth could support quality control by leveraging its training on empty and ambient droplets to distinguish high-confidence cells from unassignable or low-quality profiles (Supplementary Methods). Because Tabula Sapiens had already undergone extensive manual quality control, Pan-human Azimuth assigned the “Unassigned” label to only a small fraction of cells (0.3%). Nevertheless, marker-gene analysis supported these assignments. The most common case involved cells manually labeled as neutrophils in Tabula Sapiens but classified as “Unassigned” by Pan-human Azimuth. In these cells, the canonical neutrophil marker *FCGR3B* was largely absent, whereas it was robustly expressed in cells annotated as neutrophils by both the manual and automated approaches (Fig.3L). We observed similar patterns across additional neutrophil markers (Fig.3M) and other cell types, including cells manually annotated as macrophages but labeled as “Unassigned” by Pan-human Azimuth (Supplementary Fig.2C). These results indicate that the “Unassigned” label provides a useful quality-control output, enabling Pan-human Azimuth to flag low-confidence profiles without relying on strict or arbitrary metric-based cutoffs.

We next benchmarked its predictions against recent organism-scale annotation approaches, including the scGPT ^15^ and SCimilarity ^16^ foundation models. These comparisons are inherently challenging as foundation-model labels, which span multiple distinct studies, are neither internally consistent nor aligned with Pan-human Azimuth’s harmonized hierarchical typology. This distinction underscores that Pan-human Azimuth is not simply a marginal improvement in annotation accuracy, but a framework for standardized pan-human cell type assignment. Nonetheless, we performed systematic benchmarks to assess empirical performance.

We first evaluated annotation accuracy in the Tabula Sapiens lung atlas, focusing on epithelial populations because they are well defined and characterized by distinctive marker-gene programs. Manual Tabula Sapiens labels showed the expected canonical signatures (Supplementary Fig.3A top-left), including AT1 and AT2 markers, secretory and goblet cell programs, and ciliated-cell markers, all of which were faithfully recapitulated by Pan-human Azimuth (Supplementary Fig.3A bottom-left). In contrast, annotations from scGPT and SCimilarity were substantially noisier (Supplementary Fig.3A top-right and bottom-right). For example, SCimilarity assigned a subset of lung epithelial cells to cardiac muscle despite the absence of cardiomyocyte marker expression, consistent with over-transfer of mismatched reference labels. Similarly, goblet cells annotated by scGPT did not show coherent expression of canonical secretory markers, and lung basal cells were frequently confused with basal populations from other tissues.

We next designed a quantitative benchmark to evaluate each automated method on its own assigned labels. Since the distinct label sets returned by each method prohibited direct comparison, we asked whether the populations identified by each method were transcriptionally distinguishable. Specifically, we defined a differential-expression-based diagnostic, which we refer to as ‘DE separability’ (Supplementary Methods). For each method, we identified marker genes associated with each predicted label and then asked whether those markers were sufficient to recover the same labels using a logistic-regression classifier, with classifier training accuracy defining the metric. This approach captures whether predicted annotations are supported by coherent gene-expression programs. High DE separability indicates that labels correspond to transcriptionally distinct populations, whereas lower separability can reflect label fragmentation, over-transfer, or assignments that are not well supported by expression. Because the metric operates within each method’s own output label space, it provides a complementary way to compare annotation quality without forcing all tools into an exact shared vocabulary.

We applied this framework to all automated methods and the original manual annotations, evaluating DE separability at both the single-cell and class-wise levels. We first examined Tabula Sapiens lung epithelial cells, where Pan-human Azimuth and the manual annotations produced nearly identical separability distributions, supporting both the validity of the metric and the biological coherence of Pan-human Azimuth predictions (Supplementary Fig.3B). By contrast, SCimilarity and especially scGPT showed broader and more uniform distributions and more variable class-wise performance, consistent with fragmented or weakly supported label sets. Similar patterns were observed across multiple tissues in both immune and epithelial compartments (Supplementary Fig.4), consistent with the expected limitations of training annotation models on heterogeneous, author-defined labels.

### Scalable and unified cell type annotation with Pan-human Azimuth

We next applied Pan-human Azimuth to scBaseCamp ^25^. To our knowledge, scBaseCamp is the largest single-cell RNA-seq compendium assembled to date, containing tens of million human scRNA-seq and snRNA-seq profiles that were systematically mined from public databases using agentic workflows. Although unique in scale, the resource is largely a raw compendium of expression profiles and metadata, but lacks cell type annotations due to existing challenges in providing standardized annotations across the breadth of tissues. This makes scBaseCamp a powerful use case to evaluate whether Pan-human Azimuth can scale to a highly heterogeneous organism-scale compendium.

After restricting to non-cancer human samples, consistent with the scope of our training data, we curated a dataset of approximately 85.9 million cells spanning 11,398 SRX accessions and annotated the full collection using Pan-human Azimuth. The complete annotation procedure required approximately 13.5 hours on an NVIDIA A100 GPU and 23.5 hours on an Intel Xeon CPU. Based on the model’s built-in QC, 11% of profiles were marked as “Unassigned”; these profiles had substantially lower UMI counts than cells assigned a cell type label, with median UMI counts of 19 and 4,734, respectively. The remaining ~75 million cells were generally annotated with high confidence, with a median confidence score of 0.89, and spanned a broad range of cell types and tissues (Fig.4A–C).

**Figure 4.**
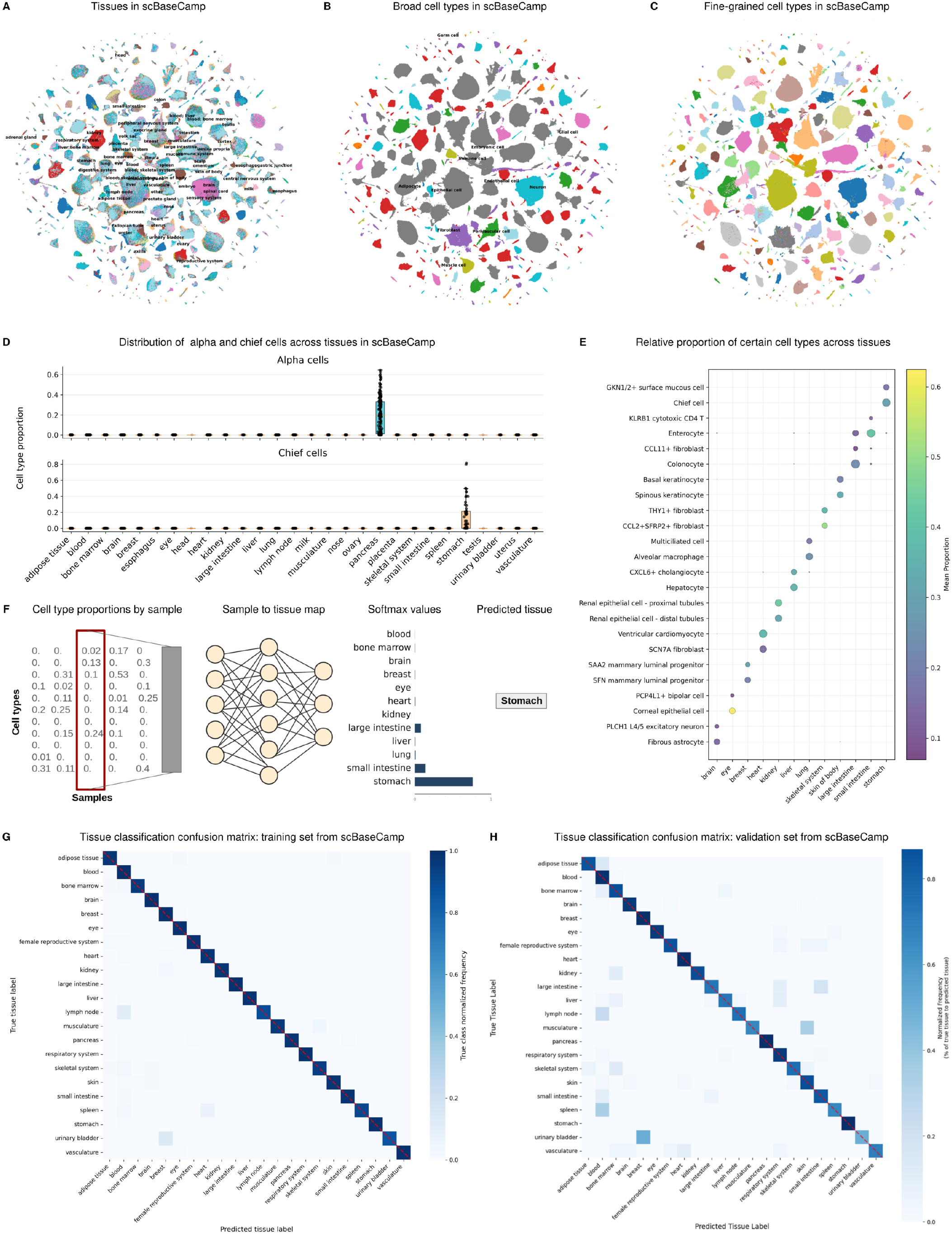
Annotating scBaseCamp with Pan-human Azimuth. (A) UMAP visualization of Pan-human Azimuth embeddings of non-cancer human data in scBaseCamp. Cells are colored by tissue metadata (B) UMAP visualization of the broad cell type annotations from Pan-human Azimuth. (C) UMAP visualization of finely resolved cell type annotations from Pan-human Azimuth. (D) The distribution of alpha and chief cells in scBaseCamp data, as shown over a broad panel of tissues. (E) Dot-plot showing tissue specificity of different cell types whose proportion is enriched across different tissues in the body (F) Schematic of a map from cell type composition to tissue, as learned with a shallow neural network with a single hidden layer. (G) Confusion matrix of tissue classification map on the training set from scBaseCamp. (H) Confusion matrix of tissue classification map thus learned on the held-out validation set.

The annotated scBaseCamp repository represents a powerful structured resource for biological discovery across more than 10,000 datasets. We first used this resource to identify cell type marker genes that generalize across datasets, reasoning that robust markers should distinguish the same annotated cell type across thousands of samples and donors. To do so, we applied TF-IDF analysis to quantify the discriminative power of each gene for distinguishing annotated cell types, then aggregated gene-importance scores across samples to identify the top 50 markers for each cell type (Supplementary Methods). Visual inspection confirmed that these markers were biologically interpretable and consistent with known cell identities, including markers for abundant populations such as *CD3D* and *CD8A* in CD8 T cells, as well as rare populations such as *FOXI1* in pulmonary ionocytes ^62,63^ and *AXL* in ASDCs ^64^. To assess robustness, we repeated the same analysis in the Tabula Sapiens atlas^23^, which was not included in scBaseCamp. Marker overlap between datasets was substantial: across cell types and over the shared panel of genes between the two databases, a median of 63% of the top 50 scBaseCamp markers were also recovered in the top 50 markers from Tabula Sapiens (Supplementary Fig.5A). Our marker lists are provided in Supplementary Table 2 and have been incorporated into the updated HuBMAP cell type and biomarker reference tables ^65^.

We also evaluated whether Pan-human Azimuth could support interpretation of diseased datasets, despite being trained primarily on healthy, non-cancer reference data (Supplementary Methods). Encouragingly, when we restricted the analysis to scBaseCamp samples with a disease label (non-cancer), marker genes derived from healthy reference data remained strongly discriminative across annotated cell types (Supplementary Fig.5B). This indicates that, in many settings, diseased cells preserve enough cell-type-specific transcriptional structure to support reliable annotation against a healthy atlas. Pan-human Azimuth can therefore provide an initial harmonized cell type scaffold for diseased datasets, enabling downstream cell type-resolved analyses to identify disease-associated transcriptional changes within each population. Although this does not replace the need for disease-specific reference atlases, particularly for cancer, which is outside the scope of Pan-human Azimuth, it supports the broader applicability of the framework beyond strictly healthy datasets.

### Cell type distributions across tissues

We next asked whether the assigned labels recovered reproducible sample-level patterns of tissue-specific cell type composition. The large number of samples in this resource allowed us to treat samples, rather than individual cells, as the unit of comparison. We generated a harmonized sample-by-cell type composition matrix that allowed us to quantify reproducible patterns of cell type abundance across tissues. Conceptually, this analysis mirrors differential expression profiling, but instead of quantifying which genes are enriched within a given cell type (with individual cells as the unit of analysis), we quantified which cell types are enriched within each tissue (with individual samples as the unit of analysis). As initial sanity checks, we examined well-established tissue-restricted cell types, for example alpha cells in the pancreas (Fig.4D top), chief cells in the stomach (Fig.4D bottom), and cardiomyocytes in the heart (Fig.4E). In each case, these cell types were robustly detected only in their expected tissues, with negligible background signal elsewhere.

We identified representative examples of tissue-enriched cell types across multiple contexts (Fig.4E), and next asked whether tissue of origin could be predicted from cell type composition alone. To our knowledge, no tissue classifier based on scRNA-seq sample composition currently exists, but the breadth of scBaseCamp provided a unique opportunity to build one. We therefore trained an MLP classifier to predict each sample’s tissue of origin using only the relative abundances of Pan-human Azimuth–assigned cell types (schematic in Fig.4F, Supplementary Methods). The classifier achieved high accuracy on both the training and held-out scBaseCamp samples (Fig.4G–H), demonstrating that Pan-human Azimuth annotations recover reproducible tissue-specific patterns of cellular composition. Performance was also strong on the independent Tabula Sapiens atlas (Supplementary Fig.5C), indicating that these compositional patterns generalize across datasets.

**Figure 5.**
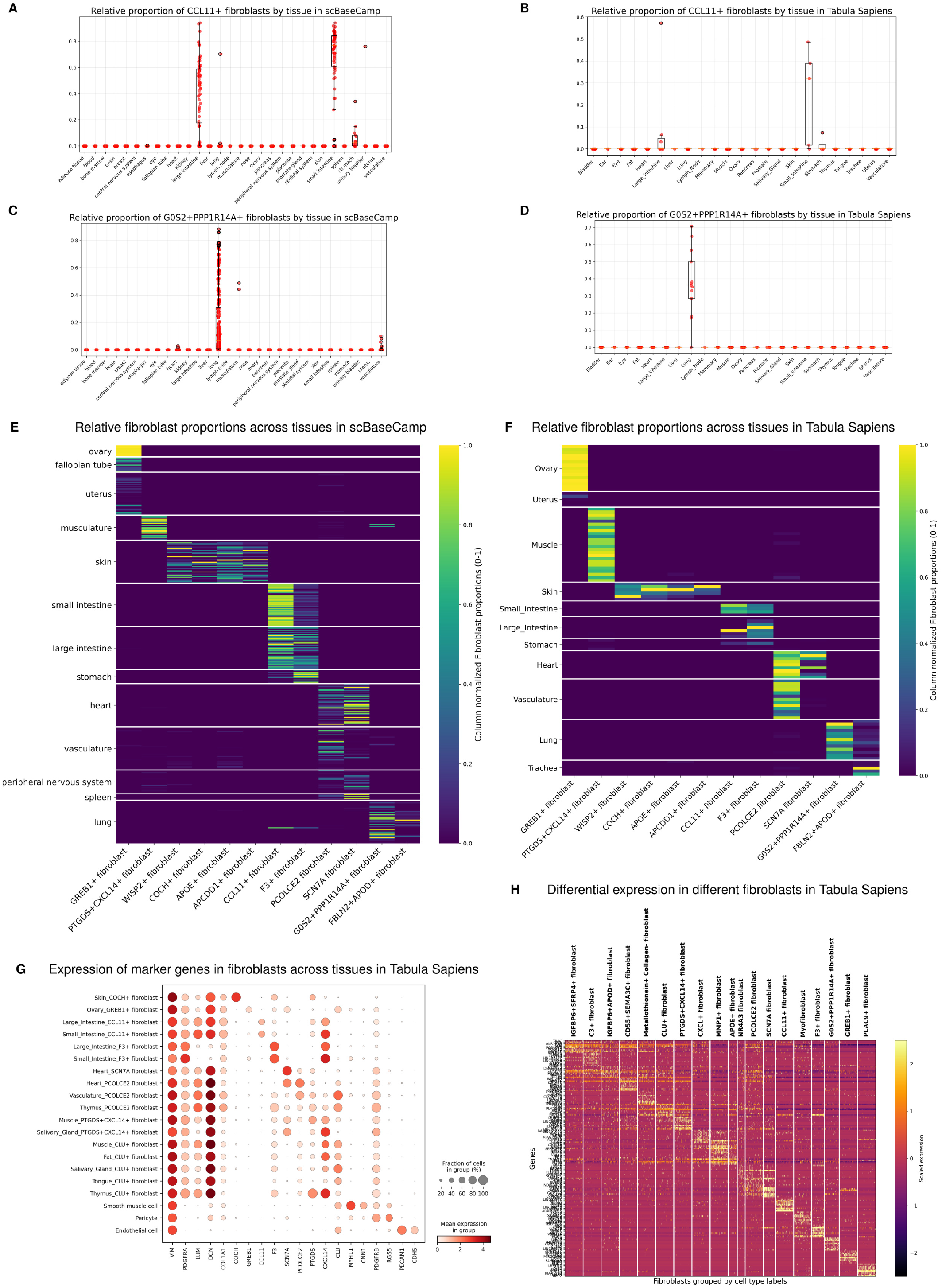
Tissue specificity of fibroblasts. (A) Relative proportions of CCL11^+^ fibroblasts across different tissues in scBaseCamp show that these fibroblasts are specialized to the gastrointestinal tract. (B) Specificity of CCL11^+^ fibroblasts to the intestines and the stomach is also observed in data from Tabula Sapiens. (C) Relative proportions of G0S2^+^ PPP1R14A^+^ fibroblasts in scBaseCamp show specificity to lung tissue. (D) Localization of G0S2^+^PPP1R14A^+^ fibroblasts to the lung is replicated in data from Tabula Sapiens. (E) Heatmap of fibroblast-state proportions across tissues in scBaseCamp, revealing distinct patterns of tissue specialization. (F) The same tissue-associated fibroblast-state patterns are recapitulated in Tabula Sapiens. (G) Dot plot showing expression of marker genes across selected tissue-associated fibroblast populations and related mesenchymal cell types, including smooth muscle cells and pericytes. Fibroblast populations are clearly distinguished from other mesenchymal lineages, and cell-type-specific marker programs are preserved across tissues despite weaker tissue-associated transcriptional variation. (H) Differential-expression analysis across major fibroblast populations demonstrates that these fine-grained annotations are supported by distinct gene-expression programs.

We performed feature-importance analysis (Supplementary Methods) to identify the cell types driving tissue classification and found that epithelial populations were among the strongest discriminators of tissue identity (Supplementary Fig.5D), consistent with their well-known tissue specificity. Building on this observation, we asked whether tissue identity could be encoded not only by highly tissue-restricted cell types, but also by more broadly distributed lineages that adopt tissue-specialized states. We focused on fibroblasts because they are present across many organs, yet are known to diversify in response to local tissue environments^66,67^. This question has been difficult to address systematically because fibroblast subtypes are often annotated separately within individual tissues, preventing direct comparison across organs. Pan-human Azimuth overcomes this limitation by applying a shared fibroblast hierarchy across all scBaseCamp datasets, enabling us to compare fibroblast state distributions across the body.

Using this framework, we found that fibroblast states span a continuum from broadly distributed to highly tissue-localized populations. Some states, including C3^+^ fibroblasts, IGFBP6^+^APOD^+^ fibroblasts, and myofibroblasts, were detected across many tissue categories (Supplementary Fig.6A-C). At the opposite end of the spectrum, several high-resolution states showed striking tissue localization. For example, CCL11^+^ fibroblasts were almost exclusively detected in the gastrointestinal tract, and G0S2^+^PPP1R14A^+^ fibroblasts exhibited the same pattern for lung samples (Fig.5A-D). Across the scBaseCamp repository, G0S2^+^PPP1R14A^+^ fibroblasts were detected in more than 60% of lung samples, comprising a median of 10% of fibroblasts in those samples. However, they were detected in fewer than 0.1% of non-lung samples, and even these included errors in the automated metadata assignment (Supplementary Fig.7C). Additional localized examples included COCH^+^ (Supplementary Fig.7A-B) and APCDD1^+^ fibroblasts, which were largely restricted to skin and cornea.

We next asked whether fibroblast state composition itself was reproducible across datasets. This provides an orthogonal validation of the annotations: just as reproducible differential-expression markers support the existence of bona fide cell types, reproducible differences in fibroblast state abundance across independent datasets would indicate that these annotations capture real tissue-associated variation rather than dataset-specific noise. Across scBaseCamp, Pan-human Azimuth-assigned fibroblast states formed a structured tissue map, with a set of distinct populations enriched in specific tissues (Fig.5E), and these compositional patterns were reproduced in the independent Tabula Sapiens atlas (Fig.5F). Thus, fibroblast state proportions vary systematically across tissues, revealing reproducible stromal composition signatures that can be resolved through harmonized cross-tissue annotation.

To confirm that these annotations reflected genuine biological states rather than over-fragmentation, we examined their transcriptional programs in the Tabula Sapiens atlas. In Tabula Sapiens, Pan-human Azimuth-defined fibroblast populations showed interpretable marker-gene programs, including expression of the genes used to define tissue-enriched states such as *CCL11, COCH, GREB1, PCOLCE2, F3*, and *CLU* (Fig.5G). More broadly, differential-expression analysis across fibroblast labels revealed distinct blocks of marker genes for each major population, demonstrating that these highly resolved annotations are supported by coherent transcriptional signatures rather than arbitrary subdivision of a continuous fibroblast compartment (Fig.5H).

Together, these analyses reveal that tissue-specialized fibroblast states are a widespread and reproducible feature of human stromal biology. While fibroblasts are known to be shaped by their tissue environment^66,67^, our results extend this idea by showing that the distribution of fibroblast states itself can encode tissue identity. We note that this pattern could only be detected by applying a uniform fibroblast hierarchy across the body, allowing stromal populations from many organs and studies to be compared directly. More broadly, these results illustrate how harmonized cross-tissue annotation can reveal organism-scale cellular organization that is difficult to resolve within individual tissue-specific analyses.

### Pan-human Azimuth extends to spatial transcriptomics data

As a final test, we evaluated whether Pan-human Azimuth could accurately annotate high-resolution spatial transcriptomic data (Fig.6A–B, Supplementary Methods). Although Pan-human Azimuth was not trained on spatial data, emerging spatial technologies such as Visium HD ^26^ measure transcriptome-wide expression at resolutions that can approximate single-cell profiles while preserving spatial coordinates. We considered Visium HD data, which provides expression measurements in 2 *µ*m bins, from a section of the kidney cortex^27^. These bins are often aggregated into larger fixed bins, for example at 8 *µ*m resolution, a step that mitigates sparsity but can blend transcripts from multiple adjacent cells and therefore violate the single-cell assumptions underlying our model. Indeed, when Panhuman Azimuth was applied to fixed 8 *µ*m bins, the large majority (~69%) of profiles were assigned the label “Unassigned” (Fig.6C and Supplementary Fig.8A). We therefore adopted a segmentation-based workflow (Supplementary Methods) in which 2 *µ*m bins are dynamically grouped according to inferred cell boundaries in order to represent single-cell profiles.

**Figure 6.**
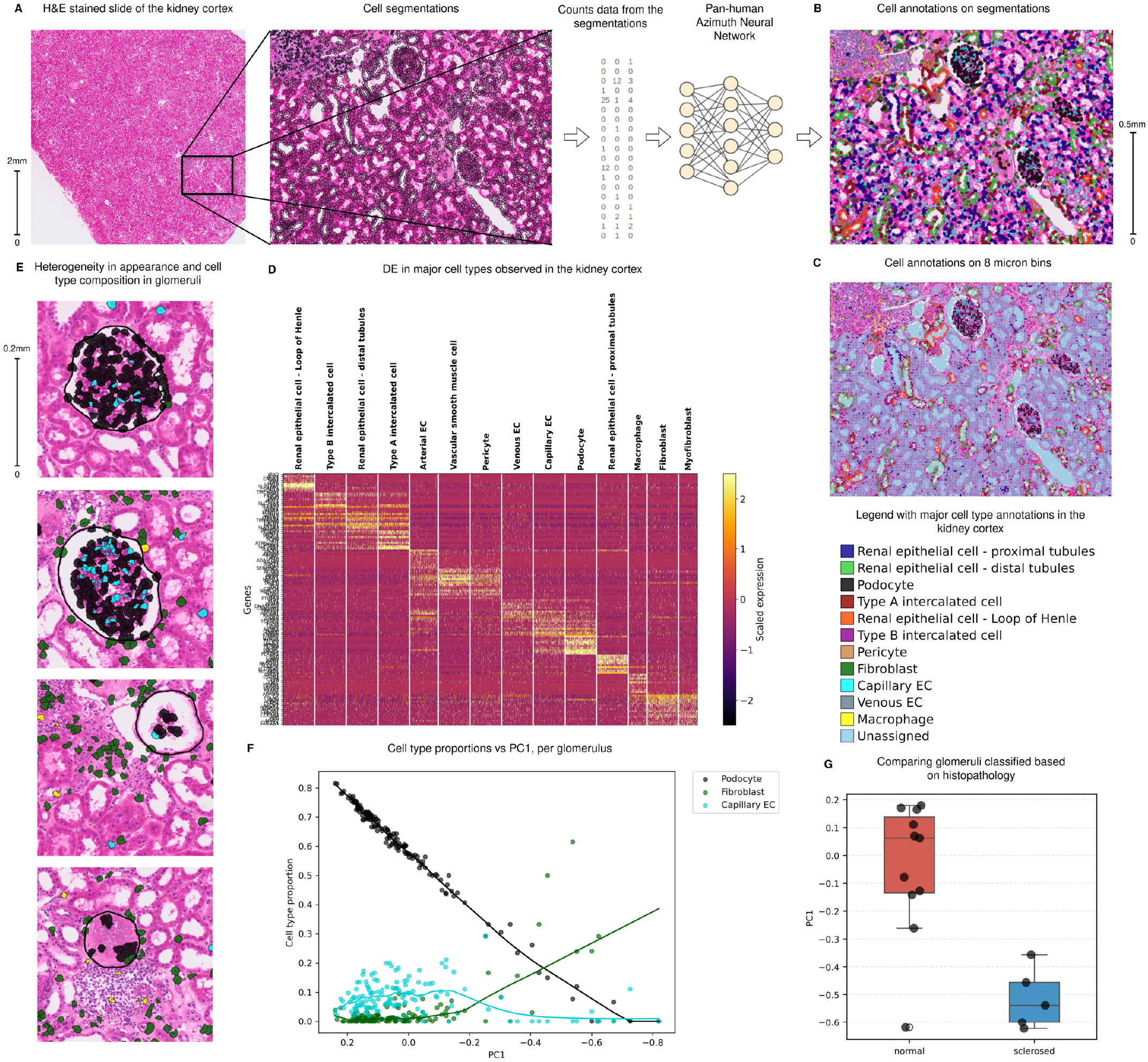
Pan-human Azimuth annotation of spatially-resolved transcriptomic data. (A) Schematic illustrating the application of Pan-human Azimuth to an H&E-stained section of human female kidney FFPE tissue profiled with Visium HD. (B) Spatial distribution of Pan-human Azimuth annotations across the kidney cortex. Although the model does not use spatial coordinates or neighborhood information, canonical cortical structures emerge from the organization of cell types assigned from molecular profiles alone. (C) Comparison with fixed 8 *µ*m bins. These spatial transcriptomic profiles provide less clearly resolved tissue structure, and most bins are assigned to the “Unassigned” category. (D) Differential-expression patterns across major annotated populations support the assigned cell identities, including kidney-specific cell types such as podocytes. (E) Four representative glomeruli identified using an independent histopathology-based segmentation model, illustrating heterogeneity in morphology and cell-type composition. (F) Principal component analysis of glomerular cell-type composition. Decreasing PC1 is associated with reduced podocyte abundance, a weaker reduction in intraglomerular capillary endothelial cells, and increased fibroblast-like populations. (G) Comparison of PC1 between glomeruli classified by an expert renal pathologist as normal or sclerosed. The observed shift in PC1 is consistent with the histopathologic classifications and supports the presence of molecular heterogeneity across glomeruli.

The segmentation-based workflow enabled us to assign annotations to ~80% of cells and accurately annotate major cell types in the kidney cortex such as podocytes, epithelial cells of the proximal and distal tubules, and intercalated cells (Fig.6B). We assessed annotation accuracy in two ways. We first looked at the differential expression of canonical markers for major cell types (Fig.6D). We observed expression of canonical markers in annotated podocytes, proximal and distal tubule epithelial cells, and Type A and Type B intercalated cells. These expression patterns matched markers we previously identified for scRNA-seq (Supplementary Table 2), affirming both the accuracy of our annotations and the ability of segmentation-based analysis of Visium HD data to return expression profiles that resemble scRNA-seq data.

We next asked whether Pan-human Azimuth, which used only the expression profile of a segmented region and not its spatial position for annotation, was nevertheless able to recover the expected spatial organization of cell types in the kidney cortex (Fig.6B, Supplementary Fig.8B). Reassuringly, annotations were highly concordant with known renal histology. For example, proximal and distal tubular epithelial cells were localized along tubular profiles throughout the cortex, consistent with their expected abundance and organization in the renal parenchyma. As expected, Type A and Type B intercalated cells were annotated in spatially distinct sections of the renal tubules, consistent with the restricted localization of intercalated cells within the distal nephron and collecting duct system.

To further assess spatial consistency of our annotations, we focused on podocytes, which are highly specialized cells in the kidney glomeruli that serve an important role in protein filtration during the formation of urine ^68,69^. To independently validate their specific localization to glomeruli, we generated glomerular masks and outlines using an existing SegFormer B5^70^ model trained on Kidney Pathology Image Segmentation (KPIs) data ^71,72^ (Supplementary Fig.8B bottom, Supplementary Methods). These masks showed near-perfect agreement with podocyte annotations from Pan-human Azimuth (Fig.6E and Supplementary Fig.8B). We additionally observed enrichment of capillary endothelial cells within the glomerular interior (Fig.6E), consistent with the dense capillary network that characterizes this compartment^69^. These results indicate that Pan-human Azimuth can recover organizational principles of the kidney microanatomy from gene expression alone.

Closer visual inspection of individual glomeruli revealed substantial heterogeneity in both morphology and cellular composition (Fig.6E). Individual glomeruli differed in apparent cross-sectional area, luminal thickness, and the regularity of their boundaries. Although some of this variation likely reflects differences in the plane of tissue sectioning, we also observed corresponding shifts in the relative abundance of podocytes, capillary endothelial cells, and fibroblast-like populations within and surrounding individual glomeruli. Together, these patterns suggested that the glomeruli captured in this section may occupy distinct functional or pathologic states.

To investigate this further, we performed principal component analysis (PCA) on glomerular cell type composition across all glomeruli in the dataset (Supplementary Methods). The first principal component (PC1) captured a clear gradient marked by progressively decreasing podocyte abundance (Fig.6F). At one extreme, glomeruli were enriched for podocytes, contained moderate to high relative proportions of capillary endothelial cells, and showed minimal representation of fibroblasts. At the opposite extreme, podocytes were nearly absent and podocyte loss was accompanied by a substantial increase in fibroblast abundance, consistent with a disrupted and fibrotic glomerular state. Glomeruli at this end of the spectrum also exhibited reduced cross-sectional area (Supplementary Fig.8C), although measurements of cross-sectional area will exhibit technical variation driven by sectioning. Collectively, these features are consistent with the interpretation that low-PC1 glomeruli represent injured or degenerating states, which are classically associated with podocyte loss, fibroblast infiltration, and structural collapse.

We note that although this dataset was generated from a sample without reported clinical kidney disease, glomerulosclerosis can be observed in healthy individuals and becomes more common with age and kidney injury^73,74^. We therefore asked whether the heterogeneity we observed across glomerular states corresponded to conventional histopathologic assessment of sclerosis. To address this, an expert renal pathologist assigned a binary classification of normal or sclerosed to a subset of glomeruli, and we compared podocyte, capillary endothelial cell, and fibroblast fractions, as well as PC1 and cross-sectional area, across these two groups. We found near-perfect concordance between our PC1 score and the predicted changes in cell type proportions with these binary classifications (Fig.6G and Supplementary Fig.8D). In addition to demonstrating agreement between binarized molecular and pathological analyses, our findings also suggest that molecular profiling may capture a more continuous spectrum of states among non-sclerosed glomeruli that appear healthy by routine histology alone.

## DISCUSSION

We introduce Pan-human Azimuth, an organism-wide framework for unified and scalable cell annotation of human single-cell and single-nucleus RNA sequencing data. Rather than requiring users to choose among tissue-specific references or reconcile inconsistent labels across studies, the model projects cells from diverse tissues into a unified annotation tree, with each cell receiving up to eight hierarchically structured labels. We demonstrate that Pan-human Azimuth exhibits robust and state-of-the-art annotation performance, can scale to annotate tens of millions of cells, and can be used to interpret both dissociated single-cell and spatially resolved datasets.

A central theme from our study is that the quality and organization of training data can be as important as model architecture or training scale. Rather than training on the largest possible collection of data, we invested in creating a consistent annotation framework: harmonizing labels across tissues, organizing them into a unified hierarchy, correcting local labeling errors, filtering low-confidence cells, and including examples of empty droplets and ambient RNA. This strategy highlights the complementarity between general-purpose foundation models and task-specific supervised models, and demonstrates that curated supervised models may be especially effective for structured and well-defined tasks.

Pan-human Azimuth provides broad biological coverage, while also pointing to clear opportunities for future expansion. For example, the current model was not designed to capture developmental stages, disease-specific states, or cancer programs. Nevertheless, our analysis of scBaseCamp suggests that many non-cancer disease datasets can still be meaningfully interpreted using a healthy reference. Cancer is likely to present a distinct challenge, since malignant cells can substantially disrupt normal molecular identity patterns and may not be interpretable using healthy reference atlases. For these contexts, disease-specific and cancer-specific atlases will be essential, and the construction of such atlases may benefit from adopting a similar practical template for data curation and task-specific training.

Particularly in single-cell genomics, an ongoing challenge for all reference models is how to incorporate new data as the field evolves. Large emerging datasets may increase model power, improve resolution within existing cell classes, reveal missing branches of the hierarchy, or define entirely new cell types and states. One option is to periodically retrain reference models from scratch, but this may become increasingly burdensome as atlases grow and as community resources are maintained by large consortia. An attractive alternative is incremental learning, in which additional training steps are focused on new datasets while preserving the structure and performance of the existing model. Developing robust strategies for incremental learning represents a promising approach for keeping organism-scale models current without requiring complete reconstruction each time new data becomes available.

Our work is focused on human datasets, but an important direction for future development is expansion to additional organisms. This would bring mouse and other model organisms ^75–77^ under the umbrella of a unified hierarchical cell annotation framework, and may benefit from the recent development of ‘universal’ or ‘cross-species’ embedding frameworks that can jointly model data across multiple species ^18,78–80^. The broad lineage levels of the existing taxonomy provide natural anchors for cross-species alignment, while finer levels can accommodate species-specific cell states, enabling principled cross-species comparisons of cell type composition and tissue specialization at a scale not currently achievable with existing tools.

Despite these challenges, organism-wide annotation models create a major opportunity for single-cell biology. By mapping tens of millions of cells across thousands of scBaseCamp samples in a single run, Pan-human Azimuth shows how a shared annotation language can enable analyses that would otherwise be difficult or impossible. For example, our discovery of reproducible, tissue-localized fibroblast states depended on the ability to compare stromal populations across the body using consistent labels. More broadly, just as the reference genome provides a common coordinate system for sequencing reads, an organism-wide cellular transcriptomic reference can provide a common language for describing cellular identity. Pan-human Azimuth aims to support this goal and to help facilitate standardized, reproducible, and integrative analyses of human cellular diversity.

## DATA AND CODE AVAILABILITY

The Pan-human Azimuth model is implemented in `panhumanpy`, a Python package that provides access to the trained model weights and supports the mapping and annotation of new datasets. The package is designed to support future model improvements to allow for subsequent updates while maintaining backwards compatibility and the ability to reproduce existing results. Installation instructions, a quick-start guide, and tutorials are available at www.satijalab.org/pan_human_azimuth.

Users can access `panhumanpy` through several interfaces. Within Python, the package provides a high-level interface that implements the default annotation workflow on AnnData objects, as well as lower-level functions that offer experienced users greater control. The package also contains a command-line interface (CLI).

R and Seurat users can access the same underlying functionality through the `AzimuthAPI` (https://github.com/satijalab/AzimuthAPI), which provides an R interface to the python package and enables the mapping of Seurat objects. The R interface also provides access to the `CloudAzimuth` function which performs inference on the cloud and does not require a local Python installation.

The `panhumanpy` source code is publicly available on GitHub (https://github.com/satijalab/panhumanpy), and the package is distributed through the Python Package Index (PyPI; https://pypi.org/project/panhumanpy/). The trained model weights are available on Zenodo ^81^, and the code used to generate the training corpus is available at https://github.com/satijalab/pha-training-data. A crosswalk between Pan-human Azimuth cell-type annotations and Cell Ontology terms is available at https://doi.org/10.48539/HBM727.TLKL.237.

## METHODS

### Curation of the training dataset

The sections below summarize the major data-processing, re-annotation, quality-scoring, and negative-example generation steps; the repository contains the full corresponding implementation details for each step.

Our training corpus integrated single-cell RNA-seq data from six Azimuth^9^ tissue references (heart, liver, pancreas, adipose, spleen, and bone marrow) ^28–46^, 21 DISCO tissue references ^11^, 123 datasets from the CZ CELLxGENE Discover portal^47,48^, 5 tissues from the GTEx ^49,50^ single-nucleus atlas (breast, heart, lung, skeletal muscle, and skin), and 215 datasets from HuBMAP ^6,7,51^. DISCO tissue references were downloaded as preprocessed Seurat objects from Zenodo. Azimuth references were built from the publicly available Azimuth reference repository (https://github.com/satijalab/azimuth-references). For CELLx-GENE, we queried the CELLxGENE Census release 2023-05-15 through the Python API, restricted to human cells marked as primary data, and exported expression matrices and associated metadata as AnnData objects organized by tissue. GTEx data were obtained from the publicly released multi-tissue single-nucleus atlas through the GTEx Portal (https://gtexportal.org/home/singleCellOverviewPage). HuBMAP datasets were retrieved from the HuBMAP Data Portal ^51^ using dataset UUIDs and downloaded as processed expression objects from the consortium asset repository. The sections below describe duplicate removal, systematic error correction, and quality scoring applied prior to model training.

#### Removal of duplicate datasets

Duplicate or redundant datasets can arise when the same samples are deposited across multiple public repositories or re-analyzed in meta-atlas efforts. In these cases, the same original cell may not have identical names or expression values after being processed through different pipelines, but should nevertheless remain extremely close to itself relative to unrelated cells from the same tissue. We therefore sought to identify cells that were nearly identical in expression space, rather than requiring exact barcode or count-matrix identity.

To identify and remove these redundancies, we compared each query dataset against the corresponding tissue-specific reference using an overlap-scoring procedure. For each dataset pair, we identified variable features in both objects, restricted the analysis to shared features, and applied log-normalization using Seurat. We then projected both datasets into a shared low-dimensional space using a random Gaussian projection matrix, with 100 dimensions by default, to enable scalable pairwise comparison while approximately preserving cell-cell distances. Nearest-neighbor relationships were computed between projected cells across datasets, and query cells whose nearest neighbor in the reference fell within a Euclidean distance threshold of 1 in the projected space were flagged as likely redundant. Datasets or cell subsets flagged during this step were excluded from downstream tissue-level integration. Duplicate removal was performed prior to reference mapping and all subsequent annotation steps.

### Error corrections and re-annotations

Initial cell type labels were generated separately for each tissue by transferring annotations from a tissue-matched reference atlas to each query dataset. This first-pass mapping was performed with anchor-based label transfer in Seurat ^53^. Reference atlases were selected on a tissue-by-tissue basis to maximize coverage and interpretability of the expected cell types. The Azimuth human adipose reference was used for adipose tissue. For lymph node and spleen, we used the CellTypist organ atlases ^8^. For brain, we used a composite whole-brain atlas derived from the Human Brain Cell Atlas v1.0 ^52^, and for bone marrow we used the BoneMarrowMap atlas^82^. DISCO references were used for the remaining cases. The HuBMAP Anatomical Structures, Cell Types, and Biomarkers working group harmonized each of the labels produced by these atlases to a single organism-wide tree.

To perform label transfer, query datasets were grouped by common tissue of origin. They were then merged, normalized, and processed with a common feature set. We identified up to 5,000 variable genes per tissue and removed features likely to introduce technical rather than biological signal, including uninformative long non-coding RNA symbols and genes with antisense suffixes. Reference anchors were computed over the first 50 dimensions of the reference reduction, and transferred labels were stored as the initial predicted cell type for each query cell.

#### Dataset-specific clustering and candidate error detection

The initial mapping step was fully automated, but could propagate systematic errors that occurred during label transfer. We therefore applied a tissue-specific correction procedure after label transfer. We used high-resolution clustering to identify local neighborhoods in which the transferred labels were internally inconsistent. Clustering was performed separately for each dataset within a tissue, so that candidate errors were evaluated relative to the structure of each original dataset. For each dataset, we constructed a nearest-neighbor graph using the PCA space and performed clustering at high resolution (resolution of 3).

To make this high-resolution clustering workflow scalable for large tissues, we used our sketch-based clustering workflow ^83^. Each tissue object was sketched with leverage-score sampling to preserve rare populations. Sketch sizes were specified separately for each tissue and source rather than as a single global fraction, so that large public compendia did not overwhelm smaller reference or source-specific datasets. We targeted 30,000-60,000 sketched cells from each major source, with lower targets for smaller sources or tissues with fewer available cells. Clusters learned on the sketched cells were then projected back to the corresponding full dataset, giving every cell in the full object a dataset-specific high-resolution cluster assignment.

For each dataset-specific cluster, we tabulated the initially transferred cell type labels. Clusters containing more than one transferred label were treated as candidates for local reannotation. Mixed labels within a compact high-resolution neighborhood could represent real biological differences not detected by unsupervised clustering, or could reflect errors in cell annotation. Within each such cluster, we identified the most abundant label as the majority cell type and all other labels as minority cell types. Within each tissue, the unique majority–minority label pairs observed across clusters were used as candidate comparisons for the pairwise cluster-level re-annotation workflow.

#### Pairwise cluster-level re-annotation

We used pairwise cluster-level re-annotation to correct local annotation errors that arose when transcriptionally similar cell types were assigned different initial labels within the same cluster. The rationale was that, if a classifier trained to distinguish two reference cell types consistently reassigns cells from one label to the other, this provides evidence that the original label reflects a mapping error rather than a distinct transcriptional population. Conversely, the classifier also allowed cells to retain their original label when their expression profiles remained better supported by the cell type assigned through label transfer.

For each candidate pair of co-clustered cell types, we trained a reference-based pairwise classifier to evaluate whether cells assigned to the minority label were better supported by the majority label, or whether the reverse was true. Marker genes were selected from the tissue reference by differential expression between the two cell types in both directions (selecting at most 100 markers in each direction based on the Wilcoxon rank-sum test). Models were fit on normalized reference expression values from cells belonging to the two compared cell types.

Models were fit using `glmnet::cv.glmnet` with a binomial likelihood and ridge penalty (`family = “binomial”`, `alpha = 0`), using the default 10-fold cross-validation to select the regularization parameter. To mitigate the effects of class imbalance (i.e. when one class contained more than 20 times as many reference cells as the other), we used downsampling of the majority class and Synthetic Minority Over-sampling Technique (SMOTE). We then applied these classifiers to the candidate cells identified for possible re-annotation in the previous section. For each cell, we computed the probability of membership in the alternative cell type using the corresponding pairwise model. If the predicted class differed from the initial transferred label at a probability threshold of 0.5, the cell was relabeled.

#### One-versus-all QC scoring

Independently of cluster structure, we trained one-versus-all logistic regression models to evaluate the transcriptional support for each assigned cell type within each tissue, stratified by data source (e.g., Azimuth, HuBMAP, CELLx-GENE). This step was designed to generate conservative annotation-quality scores rather than to relabel cells, complementing the pairwise cluster-level procedure. For each cell type, 100 marker genes, including both positive and negative markers, were selected from the reference. Predicted membership probabilities for each cell’s assigned cell type were retained as quality scores, but were not used to change labels in this step.

The one-versus-all QC scores served two purposes: filtering low-confidence annotations and prioritizing the highest-confidence cells for balanced subsampling. For each tissue, we first retained cells with one-versus-all scores of at least 0.75. Within each tissue, cells were then grouped by their corrected terminal cell type annotation, ranked by one-versus-all score, and subsampled to retain at most the top 25,000 cells per cell type. This per-cell type cap helped to prevent highly abundant populations from dominating the training corpus while preserving a large number of high-confidence examples for each label. This filtering was performed independently for each tissue before assigning hierarchical labels and concatenating tissue-specific objects into the final merged training corpus, yielding 9,665,434 high-confidence cells across 23 tissues.

#### Incorporating negative training examples

In addition to accurately annotated cells, we sought to include labeled examples of profiles that should not be assigned to any bona fide cell type. We focused on two major classes of such profiles. The first consists of empty droplets or debris, which primarily reflect ambient RNA and typically have very low RNA counts. The second consists of cell clumps, multiplets, or other mixed profiles, which can have high UMI counts but do not correspond to a single coherent cell identity and instead resemble mixtures of multiple cell types. By including negative examples from both classes, we aimed to train the model to perform quality control jointly with cell type annotation, allowing poor-quality or mixed profiles to be labeled as “Unassigned” rather than forced into an incorrect cell identity.

To collect experimentally observed low-quality profiles, we downloaded publicly available raw count matrices from twenty 10x Genomics demo datasets (https://www.10xgenomics.com/datasets), spanning diverse sample contexts including peripheral blood, jejunum, breast, brain, and lymph node. For each dataset, we identified empty droplets and debris using the inflection point of the barcode rank plot, extracting barcodes whose UMI counts fell below 50% of the threshold used for cell calling. These profiles provided empirical examples of low-count ambient RNA and droplet-associated background signal.

To complement these sparse empty droplets with higher-UMI artifacts that nevertheless lack coherent single-cell structure, we generated simulated mixture-like profiles for each tissue. Pseudobulk expression profiles were derived from Tabula Sapiens tissue objects matching the tissues represented in our training corpus. For each tissue, we computed a pseudobulk gene abundance profile that was enriched for highly expressed transcripts characteristic of ambient or mixture-derived signal. Final gene counts for each simulated profile were drawn independently from a negative binomial distribution parameterized by the pseudobulk profile and the sampled UMI total, generating 5,000 negative profiles per tissue. We reasoned that these simulated profiles would capture low-quality or non-cellular expression patterns distinct from empirically observed empty droplets alone.

In total, we generated approximately 145,000 collected and simulated negative profiles, labeled them Unassigned, and added them to the training dataset after all other curation steps were complete. During inference, cells or droplets whose expression profiles most closely resemble these negative training examples are returned with the Unassigned label, alongside calibrated confidence scores for retained annotations. This built-in quality-control output allows the model to jointly distinguish high-confidence cell identities from empty droplets, ambient RNA, and mixed or noisy profiles without relying solely on metric-based cutoffs.

### Pan-human Azimuth Neural Network

#### Training data labels

To capture the nested structure of cell type identity across varying levels of resolution and to ensure interpretable model outputs across different contexts, we framed cell type annotation as a hierarchical classification problem. For example, we wanted the model to annotate a specialized immune cell such as ‘GZMK+IL7R+ CD8 T cell’ with labels capturing the cell’s identity at multiple levels of granularity; therefore, we labeled it as ‘Immune cell|Lymphoid cell|T/NK cell|T cell|CD8 T cell|Memory CD8 T cell|GZMK CD8 T cell|GZMK+IL7R+ CD8 T cell’.

Because cell type annotation at a fine level of resolution is generally more difficult than annotation at a broader level, we split the multi-level cell type labels using a cumulative split. As shown in Fig.2C, this means that the label ‘Immune cell|Lymphoid cell|T/NK cell|T cell|CD8 T cell|Memory CD8 T cell|GZMK CD8 T cell|GZMK+IL7R+ CD8 T cell’ is split into ‘Immune cell’, ‘Immune cell|Lymphoid cell’, ‘Immune cell|Lymphoid cell|T/NK cell’, and so on for the different classification heads. This formulation allows each classification head to learn a progressively more specific decision boundary while preserving the broader lineage context inherited from preceding levels. This scheme, as opposed to splitting the label into ‘Immune cell’, ‘Lymphoid cell’, ‘T/NK cell’, and so on, ensures that annotation complexity increases with depth.

The number of classes obtained at each level was 13 (level 1), 91 (level 2), 209 (level 3) up to 382 (level 8).

#### Neural Network Architecture

The Pan-human Azimuth neural network (Fig.2A,C) is designed to perform hierarchical cell type classification, producing annotations at multiple levels of biological resolution simultaneously, while encouraging consistency across levels such that fine-grained predictions remain coherent with their parent categories. The architecture achieves this through two main components: an embedding module that compresses sparse single-cell gene expression profiles into a compact representation, followed by a nested sequence of hierarchical classification heads that decode this representation into multi-level cell type annotations, each one also conditioned on the predictions of all preceding levels.

The embedding module uses four fully connected layers to compress gene-expression profiles across 5,055 genes into a 128-dimensional latent representation. It follows the architecture 5,055 → 1,024 → 512 → 256 → 128 and comprises approximately 5.87 million parameters. This compact representation serves as the shared input to all downstream classification modules, and also provides a unified embedding coordinate system across datasets that can be used to generate two-dimensional visualizations.

The 128-dimensional embedding is then passed to a nested sequence of eight hierarchical classification heads (Levels 1–8), each predicting cell type labels at a successively finer level of ontological resolution. The number of output classes increases monotonically with depth, from 13 classes at Level 1 to 382 classes at Level 8. Each classification head follows a consistent structure: the 128-dimensional embedding is concatenated with compressed representations of all preceding levels’ softmax outputs, each passed through a small dense projection layer, and the resulting concatenated vector is passed through a dense layer followed by dropout before producing the level-specific softmax output. The hidden layer dimensions of each classification head scale with the complexity of the level, ranging from 32 units at Level 1 to 256 units at Levels 4–8. The growing concatenated input at each level (131-dimensional at Level 2, increasing to 523-dimensional at Level 8) is consistent with the increasing complexity of the classification problem at finer levels of resolution, and also ensures that each classifier has full access to the hierarchy of preceding predictions, thus directly encouraging hierarchical consistency through architectural design. The total number of trainable parameters is 6,981,993. A detailed summary of the neural network architecture, as obtained through `keras model.summary()`, is provided in Supplementary Table 1.

Cells in the training dataset are annotated up to different maximum resolutions with the overall maximum being level 8. For example, a GZMK+IL7R+ CD8 T cell has a maximum depth of 8 as it is labeled as ‘Immune cell|Lymphoid cell|T/NK cell|T cell|CD8 T cell|Memory CD8 T cell|GZMK CD8 T cell|GZMK+IL7R+ CD8 T cell’, whereas a club cell only has a maximum depth of 3 as it is labeled as ‘Epithelial cell|Epithelial cell of lung|Club cell’. In order to bring cells with varying maximum depth under the purview of the same hierarchical classification scheme, we pad any absent levels with blank labels up to level 8. So the full hierarchical label for a club cell in our dataset is ‘Epithelial cell|Epithelial cell of lung|Club cell|||||’.

#### Model loss function

We used focal loss to reduce the contribution of easily classified cells and maintain training emphasis on difficult distinctions, including rare, closely related, or fine-grained cell types. The outputs from each classification MLP were evaluated using a focal loss function, and the overall neural network was optimized with respect to a weighted aggregate of the losses computed at individual hierarchical levels. Focal loss is equivalent to cross-entropy loss supplemented with a focal factor:

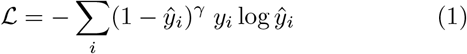

This expression defines the focal loss for one example, where the sum over *i* runs over all classes. Here, *ŷ*_*i*_ denotes the predicted softmax probability for class *i*, while *y*_*i*_ denotes the ground-truth one-hot encoding of the cell label. We set the focal parameter *γ* to 2. The focal factor (1 *− ŷ*_*i*_)^*γ*^ emphasizes difficult examples, thereby preventing training from being dominated by easier examples, such as highly abundant or transcriptionally distinct cell types.

The losses at each hierarchical level were aggregated into an overall loss function, with each level assigned a weight equal to the number of classes at that level. This weighting scheme encouraged the model to learn high-resolution cell type variation and reduced the tendency for training to saturate at shallower hierarchical levels.

#### Training

We split our final training dataset of ~9.8 million cells into training:validation:test splits with a 7:1:2 ratio. As is typical for scRNA-seq datasets, there is significant class imbalance, with cell type frequencies ranging from highly abundant to rare. To enable the model to learn rare cell type signatures effectively, we employed stratified sampling to distribute cells across splits and verify that fair class distributions are achieved by computing the relative entropy between class weights in each split and the full dataset.

Training was conducted with batch sizes of 256 using Keras 3.7.0 with TensorFlow 2.17.0 backend. We applied L2 regularization (*λ* = 0.01) to the network overall and dropout (rate = 0.2) to the dense layers in the classification modules. We used the Adam optimizer with learning rate decay on plateau at the default initial value, and with an early stopping callback monitoring validation accuracy. Optimal validation accuracy was achieved at 38 epochs. Training was performed on an NVIDIA A100-PCIE-40GB GPU, with NVIDIA Driver Version 530.30.02 and CUDA Toolkit Version 12.1.

We used classification accuracy on the ‘cumulative’-split labels at each hierarchical level as the training monitor, as this provided the most direct measure of progress during classification training. We also computed Cohen’s kappa as a metric to account for class imbalance in the dataset (Supplementary Fig.1B). As not all lineages include eight hierarchical levels, accuracy at deeper levels can be inflated by successful prediction of ‘blank’ labels. Therefore, to report final model performance, we calculated precision, recall, and F1 scores based on non-blank labels. Model performance measured in this manner on the held-out test set is shown in Fig.2B.

#### Confidence calibration

Pan-human Azimuth is equipped to provide confidence measures for cell type predictions at each hierarchical level. The final layer of each classification head outputs an apparent probability distribution via the softmax function. However, softmax outputs often do not align with true prediction accuracy in general, and deep neural networks are often overconfident^56^. Model calibration is a technique that aims to address this problem by aligning prediction confidence with observed accuracy without affecting the predictions themselves. Calibration is particularly important in our context, as users can employ confidence scores as a filtering mechanism to exclude uncertain annotations from downstream analyses.

There are several parametric and non-parametric methods in the model calibration literature^56^. We calibrate Pan-human Azimuth using a two-parameter entropy-informed temperature scaling model at each hierarchical level (Fig.2A). Temperature scaling employs a scaled softmax function to map uncalibrated logit outputs from the model to calibrated prediction probabilities.

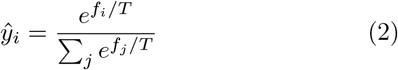

*ŷ*_*i*_ denotes the probability outputs from the scaled softmax layer, *f*_*i*_ are the logit outputs from the model with *i* ranging over all classes. The temperature *T* is a learnable parameter. In its simplest version, temperature scaling is a one-parameter model where *T* acts as the universal scaling parameter. We used a two-parameter model wherein we define *T* as a function of the entropy of the logits vector to enable it to better adapt to uncertain predictions. Thus, in our case,

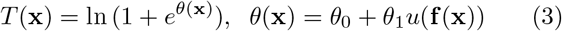

where **x** is the input gene expression vector, and *u*(**f**) is the Shannon entropy of the softmax of **f**. The scaled softmax layer is a monotonic map and thus the predictions themselves are unaffected by the calibration model.

The calibration models were trained on the validation split of the main dataset, with half of the test split reserved for validating the calibrators and the other half held out as the calibration test set. Each of these sets comprised approximately one million cells. As is standard practice, the calibration models at each level were optimized with respect to a cross-entropy loss function. Training was conducted on the GPU described above using batch sizes of 256, learning rate decay on plateau from the standard initial rate, and an early-stopping callback monitoring validation loss.

We evaluated calibration using expected calibration error (ECE) and visualized calibration performance with reliability plots. Comparing predicted labels and associated probabilities with ground-truth labels at the maximum available resolution for each cell in the calibration test dataset, we observed a post-calibration ECE of 0.0044, as shown in the reliability plot in Fig.2E. Reliability plots for all hierarchical levels are provided in Supplementary Fig.1C. In addition to reducing calibration error, the temperature-scaling models help ensure that confidence values assigned by our cell annotation tool are comparable across datasets.

### Pan-human Azimuth software interface and outputs

Pan-human Azimuth can be applied through both Python and R interfaces. The Python implementation is provided through `panhumanpy`, while the R interface is available through `CloudAzimuth`. Here, we summarize the expected inputs and primary outputs of these implementations. For operational details, we refer readers to the tutorial notebook in the GitHub repository https://github.com/satijalab/panhumanpy and to the R vignette available at https://satijalab.org/pan_human_azimuth/.

The primary input to `panhumanpy` is an AnnData object ^84^ or an h5ad filepath. `CloudAzimuth` accepts Seurat objects ^9,53,83^. In both implementations, the model returns hierarchical cell type annotations and associated confidence values, which are added to the metadata of the input object.

#### Annotation outputs

The primary outputs of the neural network are hierarchical cell type annotations and the associated confidence values. While the modular code in `panhumanpy` enables users to access all neural network outputs in detail, the important output categories directly accessible in Python are as follows. Default outputs vary slightly in the R interface.

- `level_1_labels`: the shallowest cell type labels for each cell.
- `final_level_labels`: the highest-resolution cell type labels available for each cell. The final annotation
- depth can vary across cells.
- `final_level_confidence`: the calibrated confidence value corresponding to each final-level label.
- `full_consistent_hierarchy`: a boolean indicating whether the predictions from the different classification heads are hierarchically consistent for a given cell. We describe this in more detail in a separate section below.

Users are provided a complete hierarchical label including all levels, and they can also toggle a parameter to access labels for each hierarchical level separately. The `CloudAzimuth` function in R returns three additional metadata categories by default: `azimuth_broad`, `azimuth_medium`, and `azimuth_fine`, which are optional in Python pipelines. These provide standardized cell type labels at three fixed resolutions, derived from our interpretation of the hierarchical outputs. These refined labels are particularly useful when users prefer pre-selected annotations at consistent resolutions without manually parsing the full eight-level hierarchy, especially because different cells can be annotated to different depths. We describe the label refinement algorithm in a separate section below.

#### Hierarchical consistency

Although each classification head is trained with hierarchical information, the outputs from different heads are still generated as separate predictions. As a result, a small number of cells can receive predictions that are individually plausible but mutually inconsistent across hierarchical levels. For example, a cell may be classified as a T/NK cell at one level, as an innate lymphoid cell at the next level, and then as a CD8 T cell at deeper levels.

To identify these cases, we report a metadata field, `full_consistent_hierarchy`, indicating whether the predictions from all hierarchical levels are mutually consistent for each cell. This field allows users to filter out cells with inconsistent hierarchical predictions. These cases typically correspond to expression profiles that are ambiguous to the model and represent a small proportion of quality-controlled datasets.

#### Label refinement

Raw hierarchical predictions from Pan-human Azimuth can also be post-processed into standardized labels at three fixed resolutions: broad, medium, and fine. Because the model can assign labels across up to eight hierarchical levels, and because different lineages terminate at different maximum depths, the same hierarchy level does not always correspond to the same biological resolution across cell types. The refinement step provides a simple way to collapse these variable-depth predictions into consistent broad, medium, or fine-grained annotations that are easier for users to interpret when comparing across cells, tissues, and datasets.

Label refinement is performed using predefined mapping tables distributed with `panhumanpy` and available in the GitHub repository. These tables were developed as part of the HuBMAP Anatomical Structures, Cell Types, and Biomarkers working group, which defined, for each lineage, the hierarchical depths corresponding to broad, medium, and fine annotation resolutions. Each table maps recognized hierarchical paths to curated reference annotations at the selected resolution.

For cells with consistent hierarchical predictions, the algorithm first identifies the deepest predicted hierarchical path for that cell and looks for an exact match in the appropriate mapping table. If this path is present, the corresponding broad, medium, or fine annotation is returned directly. If the model prediction stops at a shallower point in the hierarchy, the algorithm instead searches the mapping table for all recognized deeper paths that begin with the predicted prefix. Among these valid descendant paths, it selects the one with the highest corresponding model confidence and returns the annotation associated with that path at the requested resolution. Thus, label refinement converts variable-depth hierarchical predictions into standardized broad, medium, or fine labels while restricting outputs to recognized paths in the curated hierarchy.

### Tabula Sapiens v2 data analysis

Tabula Sapiens v2 data were downloaded from the public Figshare release ^85^ of the Tabula Sapiens Consortium (https://figshare.com/articles/dataset/Tabula_Sapiens_v2/27921984). This release contains the Tabula Sapiens 2.0 atlas, comprising more than 1.1 million cells from 28 tissues across 24 human donors, including nine new donors and four new tissues added to the original Tabula Sapiens 1.0 dataset. We downloaded the processed expression objects provided with the release and used the accompanying cell- and sample-level metadata for tissue, donor, and annotation information. We processed all cells with Pan-human Azimuth using default settings in the `panhumanpy` package described above.

We describe minor implementation details for individual visualization panels in Figure 3 below. We filtered out cells with hierarchically inconsistent annotations, or final-level confidence less than 0.75, for downstream analyses in Fig.3E-K. Comparisons between manual annotations and Pan-human Azimuth annotations for lung epithelial cells in the Tabula Sapiens dataset (Fig.3E) were performed using cells assigned to ‘Epithelial cell’ by Pan-human Azimuth. For ease of visualization, we removed cell labels with less than 50 examples in either the manual or Azimuth annotation sets in panels E-G. For Fig.3M, we collected the set of top 10 DE genes for all myeloid cell types in the dataset and then plotted the expression data from cells manually classified as ‘Neutrophils’, but flagged as ‘Unassigned’ by Pan-human Azimuth.

#### Assessing transcriptomic support for annotations

Tabula Sapiens v2 data were processed with two foundation models that support reference mapping. First, we ran SCimilarity ^16^ following the procedure for unconstrained annotation in the “Annotating cell types” tutorial provided in the SCimilarity documentation (https://genentech.github.io/scimilarity/notebooks/cell_annotation_tutorial.html). Second, we ran scGPT ^15^ following the tutorial for human reference mapping (https://github.com/bowang-lab/scGPT/blob/main/tutorials/Tutorial_Reference_Mapping.ipynb). All tools were run with their default parameters.

Directly comparing annotation accuracy across automated tools is challenging because different methods use different label vocabularies, annotation depths, and cell type ontologies. A prediction that appears discordant by exact label matching may reflect a difference in naming or granularity rather than an incorrect biological assignment. We therefore sought to evaluate a complementary property of each annotation set that does not require labels from different tools to be matched to one another: whether the labels assigned by a given method define transcriptionally coherent and separable populations.

We refer to this diagnostic as *DE separability*. The rationale is that, if an annotation method identifies biologically meaningful cell populations, then those labels should exhibit high in-sample separability, and be supported by differentially expressed genes. Consequently, the assigned cell type labels should be recoverable from expression of these marker genes using a simple classifier. Conversely, if a method produces labels that are noisy, redundant, or unnecessarily fragmented, the resulting groups may not be clearly distinguishable by their own marker genes. This analysis therefore measures the internal transcriptional consistency of each annotation scheme, rather than exact agreement with a single reference vocabulary.

For each benchmarking dataset and each annotation method, we considered the set of labels returned by that method, denoted *S*. We then performed differential expression analysis using those labels and identified the top 10 marker genes for each label. The union of these marker sets across all labels defined a method-specific marker gene set, denoted *G*_*S*_. We next trained a multi-class logistic regression model to predict the labels *S* using only expression values for genes in *G*_*S*_. The resulting training accuracy was used as the DE separability score. In this framework, high separability indicates that the method’s labels are well supported by marker-gene expression, whereas low separability indicates that the labels are difficult to distinguish even using the genes selected as most differentially expressed for those same labels.

We used logistic regression deliberately because it provides a simple linear test of marker-gene separability. More flexible nonlinear classifiers could potentially recover labels from complex expression patterns, but would make the metric more dependent on classifier architecture, optimization, and hyperparameter choices. By using an unregularized linear model, the analysis asks a more direct question: whether the annotation labels correspond to groups that are separable by differential expression structure.

We applied this procedure independently to manual annotations and to annotations generated by Pan-human Azimuth, scGPT, and SCimilarity. For each annotation set, marker genes were selected separately, logistic regression models were trained separately, and DE separability was evaluated with respect to that method’s own labels. Thus, the analysis does not require harmonizing labels across methods and does not assume that any one annotation vocabulary is the ground truth. The resulting DE heatmaps for lung epithelial cells from Tabula Sapiens are shown in Supplementary Fig.3A.

After fitting an independent logistic regression model for each annotation method, we examined both the overall training accuracy and the per-class accuracy distribution. Right-shifted per-class accuracy distributions indicate that most labels are strongly supported by marker-gene expression, whereas left-shifted distributions indicate labels that are poorly separable, potentially reflecting ambiguous annotations, redundant labels, or fragmentation of transcriptionally similar populations. These distributions are shown in Supplementary Fig.3B-C and can be compared with the corresponding DE heatmaps in Supplementary Fig.3A.

DE separability analyses were performed on cells with hierarchically consistent annotations that passed a minimum confidence threshold of 0.5. Multi-class logistic regression models were trained using an `sklearn`-based implementation without regularization, with the maximum number of iterations set sufficiently high to allow training accuracy to stabilize. For tissues with sufficient coverage from new donors in Tabula Sapiens v2, including spleen and stomach, comparisons were restricted to these unseen samples to avoid overlap with training data for the automated methods. For tissues with limited new data in terms of volume or donor diversity, including lung and liver, all available samples were used. Although this latter setting does not provide a strict test of generalization to unseen donors, it still provides a controlled comparison of the transcriptional separability of the annotation schemes produced by each method on the same cells.

### scBaseCamp analysis

scBaseCamp data, now distributed as part of the Arc Virtual Cell Atlas under the name scBaseCount, were accessed from the public Arc Institute data release via the Google Cloud Marketplace bucket gs://arc-institute-virtual-cell-atlas^25^. We downloaded uniformly processed scBaseCamp expression objects and associated metadata for all human datasets. We removed cancer datasets as indicated by sample metadata, but otherwise kept all healthy and diseased samples from all tissues. We processed all cells with Pan-human Azimuth using default settings in the `panhumanpy` package described above. We filtered out cells with ‘Unassigned’ labels, or hierarchically inconsistent annotations, or confidence scores below 0.75 (referred to as ‘filtered data’) for downstream analysis.

#### TF-IDF analysis to identify marker genes

The large number of samples in scBaseCamp provided an opportunity to identify marker genes that are not only enriched in a given cell type, but are consistently informative across many independent biological samples and studies. We therefore used a TF-IDF-based analysis to score genes by their relative importance to each cell type across a large collection of sample-cell type pseudobulks. In this framework, genes are treated as terms, each sample-cell type pseudobulk is treated as a document, and the full collection of pseudobulks is treated as the corpus.

The TF-IDF algorithm provides a measure of the relative importance of a term to a document within a collection of documents. For a term *t* appearing in a document *d* with frequency *f*_*t,d*_, the term-frequency component is defined as

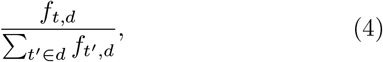

which measures the relative abundance of term *t* within document *d*. However, term frequency alone does not distinguish terms that are specific to one document from terms that are broadly present across the corpus. Specificity is captured by the inverse document-frequency component,

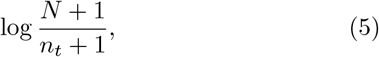

where *N* is the total number of documents in the corpus and *n*_*t*_ is the number of documents in which term *t* appears. We add a +1 to both the numerator and the denominator for smoothing. The product of these two terms gives the TF-IDF score for term *t* in document *d*.

This formulation translates naturally to marker gene identification. A gene receives a high TF-IDF score for a sample-cell type pseudobulk if it is highly expressed in that pseudobulk but is not broadly expressed across many other sample-cell type combinations. We performed pseudobulking on the filtered data using unique sample-cell type identities, with `SRX_accession` from scBaseCamp used as the sample identity and the final-level label from Pan-human Azimuth used as the cell type identity. We retained sample-cell type pairs containing at least 10 cells. This produced a pseudobulked dataset derived from approximately 46.4 million individual cells, spanning 10,639 samples, 348 cell types, and 155,399 unique sample-cell type pairs. scBaseCamp provides human gene expression data across a panel of 36,601 genes.

For the term-frequency component, we used log1p-transformed counts normalized to 10,000 per pseudobulk directly in place of the relative-frequency expression in Eq. 4. In natural language applications, an overall additive constant is often included in the IDF term to avoid eliminating ubiquitous words entirely. In our analysis, we used the basic inverse document-frequency definition in Eq. 5, which strongly downweights genes observed broadly across many sample-cell type pairs and thereby helps exclude housekeeping genes from the resulting marker lists. Finally, for each cell type, we averaged TF-IDF scores for each gene across all samples containing that cell type and retained the top 50 genes as candidate markers. We performed the same analysis on the Tabula Sapiens dataset after applying the same quality filters, using donor-tissue identities as the sample identity.

#### Pan-human Azimuth annotations in diseased and healthy samples

We next used the TF-IDF framework to evaluate whether Pan-human Azimuth annotations were associated with consistent marker-gene programs across healthy and diseased samples. This analysis leveraged the breadth of scBaseCamp to ask whether cell type annotations remained transcriptionally coherent across biological conditions, rather than being driven primarily by disease-specific transcriptional states.

Using the `disease` metadata from the 10,639 samples retained for TF-IDF analysis, we manually grouped samples into healthy and diseased sets. Samples annotated with terms such as `normal tissue`, `healthy individual`, or related descriptions were assigned to the healthy set, while samples with disease-associated metadata were assigned to the diseased set. Ambiguous samples were excluded from this comparison. This yielded a healthy set of 18,103 unique sample-cell type identities spanning 1,143 SRX accessions, and a diseased set of 81,040 sample-cell type identities spanning 4,905 SRX accessions. We then applied the same TF-IDF procedure independently to the healthy and diseased sets and calculated the overlap between the resulting marker gene sets for each cell type shared across both conditions.

#### Learning a tissue classification map

To test whether sample-level cell type composition was sufficient to recover tissue identity, we trained a tissue classification model using annotated scBaseCamp samples, as schematized in Fig.4F. We started with the filtered scBaseCamp data, but first harmonized tissue labels because the original metadata contained repeated, overlapping, or ambiguous terms. Closely related labels, such as ‘colon’ and ‘large intestine’, were merged into a common tissue label when appropriate. Samples with ambiguous tissue labels, such as ‘blood; bone marrow’, ‘blood; liver’, ‘mucosal/epithelial tissue’, or ‘other’, were excluded. As a final filtering step, we removed tissue labels represented by fewer than 20 samples.

This produced a dataset containing 8,489 samples across 22 tissue labels and 374 cell types. Samples were split into training and validation sets, with approximately 10% of samples from each tissue reserved for validation. Each sample was represented as a vector containing the relative proportion of each annotated cell type in that sample. This vector was used as input to a neural network with a single hidden layer of 160 nodes, trained to predict one of the 22 tissue labels. The network was regularized with dropout at a rate of 0.15 and trained for 100 epochs using batches of 32 samples. Optimization was performed using the Adam optimizer in Keras with `ReduceLROnPlateau` monitoring validation loss, and the best model state was retained using an `EarlyStopping` callback monitoring validation accuracy. The resulting model was evaluated on Tabula Sapiens data processed with the same quality filters. Results for tissues shared between scBaseCamp and Tabula Sapiens are shown as a confusion matrix in Supplementary Fig.5C.

#### Feature attribution for tissue classification

To identify the cell types driving tissue classification, we applied Integrated Gradients^86^ to the trained tissue classifier using the `alibi` implementation ^87^, with the Gauss-Legendre quadrature method and 100 integration steps. Integrated Gradients attributes a model’s prediction to its input features by accumulating gradients along a path from a baseline input to the sample of interest. We used the mean cell type composition vector across all validation samples as the baseline, so that attributions reflect the features distinguishing a given tissue from an average sample rather than from an empty composition. For each of the tissue labels, we computed attributions for all validation samples of that tissue with respect to its own class, and averaged the resulting attribution scores across samples to obtain a single cell type-by-tissue attribution matrix. Positive scores indicate cell types whose abundance supports assignment to that tissue, while negative scores indicate cell types whose abundance argues against it. The resulting attributions are shown in Supplementary Fig.5D.

### Analyses on spatial transcriptomic data

The analyses in this section were based on a publicly available Visium HD dataset from a human female kidney ^27^, downloaded from https://www.10xgenomics.com/datasets/visium-hd-cytassist-gene-expression-libraries-human-kidney-ffpe. Visium HD measures spatial transcriptomes at 2 *µ*m resolution, but these measurements can be aggregated into larger analysis units in multiple ways. We considered two such representations: fixed 8 *µ*m bins, which provide a regular grid-based aggregation of the underlying 2 *µ*m measurements, and segmentation-based bins, in which transcripts are aggregated according to inferred cell boundaries.

Pan-human Azimuth annotations were generated from gene expression profiles alone, as the model architecture does not use spatial coordinates or spatial neighborhood information. The spatial data included more than 666,000 fixed 8 *µ*m bins and approximately 148,000 segmentation-derived cell regions. The latter were generated using a custom implementation of StarDist ^88,89^ in the Space Ranger pipeline ^90^. Both representations were loaded and analyzed using Seurat v5.4.0 ^83^. After mapping with `panhumanpy`, we retained cells with hierarchically consistent labels and with confidence score thresholds exceeding 0.5. After comparing results (Fig.6B-C and Supplementary Fig.8A), we used the segmentation-based representation for all subsequent analyses.

#### Identifying glomeruli using a segmentation model

Glomerulus segmentations were obtained using a publicly available SegFormer B5^70^ model ^72^ trained on kidney pathology image segmentation data^71^. We applied the model to a high-resolution image of the slide with dimensions 43,999 *×* 32,674 pixels, read using the `tifffile` package. We tiled the slide into overlapping 3,000 *×* 3,000 pixel patches with a stride of 2,250 pixels along each axis. The segmentation model was applied independently to each patch to obtain glomerulus masks, and patch-level masks were then stitched back into whole-slide coordinates using a logical OR operation. The use of overlapping patches reduced the chance of missing glomeruli that were split across patch boundaries.

#### Cell type composition of glomeruli

Cells were assigned to glomeruli based on whether their spatial coordinates fell within the boundaries of each glomerulus mask. We then constructed a glomerulus-by-cell type matrix containing the fraction of cells assigned to each cell type within each glomerulus and used this matrix to analyze variation in glomerular cell type composition. PCA was performed on the glomerulus-level cell type fraction matrix with the 25 most abundant annotated cell types. The digitized image of the H&E-stained slide was separately analyzed for histopathological abnormalities. Glomeruli were annotated either as sclerosed (with scarring) or normal with no histopathological abnormality.

## ACKNOWLEDGEMENTS

We thank all the members of the Satija Lab for thoughtful discussions related to this work, and for their feedback on the software and the manuscript, and Ady Zhang for her assistance in evaluating alternative annotation strategies. We thank members of the HuBMAP consortium and the HuBMAP HIVE including Jonathan Silverstein, Philip Blood, Nils Gehlenborg, Pinaki Sarder, and Matt Ruffalo. We acknowledge the authors of the external datasets used in this study for making their valuable resources publicly available. This research was supported by grants from the NIH HuBMAP program and the NIH Office of the Director / Common Fund including 3OT2OD033760, OT2OD033756, U54DK134301, OT2OD026671, OT2OD030545, R03OD039970, and the Cellular Senescence Network (SenNet) Consortium through the Consortium Organization and Data Coordinating Center (CODCC) under award U24CA268108.

## AUTHOR CONTRIBUTIONS

R.S., S.S., G.M., Z.L., and K.B. conceived the research project. Data collection and curation were performed by Z.L. and G.M., with assistance from B.Z. S.S. led the conception and development of machine learning and subsequent computational analyses, with assistance from Z.L. Spatial transcriptomic analyses were performed by A.S. and S.S., under the supervision of R.S. Software development was led by S.S., with assistance from D.C., A.S., and Z.L. Histopathological examination was performed by J.P.G and S.J. Cell type ontology mapping was performed by N.V., A.P.-B., A.B., D.O.-S., and K.B., with review by S.S. and R.S. S.S. and R.S. wrote the manuscript with input from all authors.

---

**Supplementary Figure 1.**
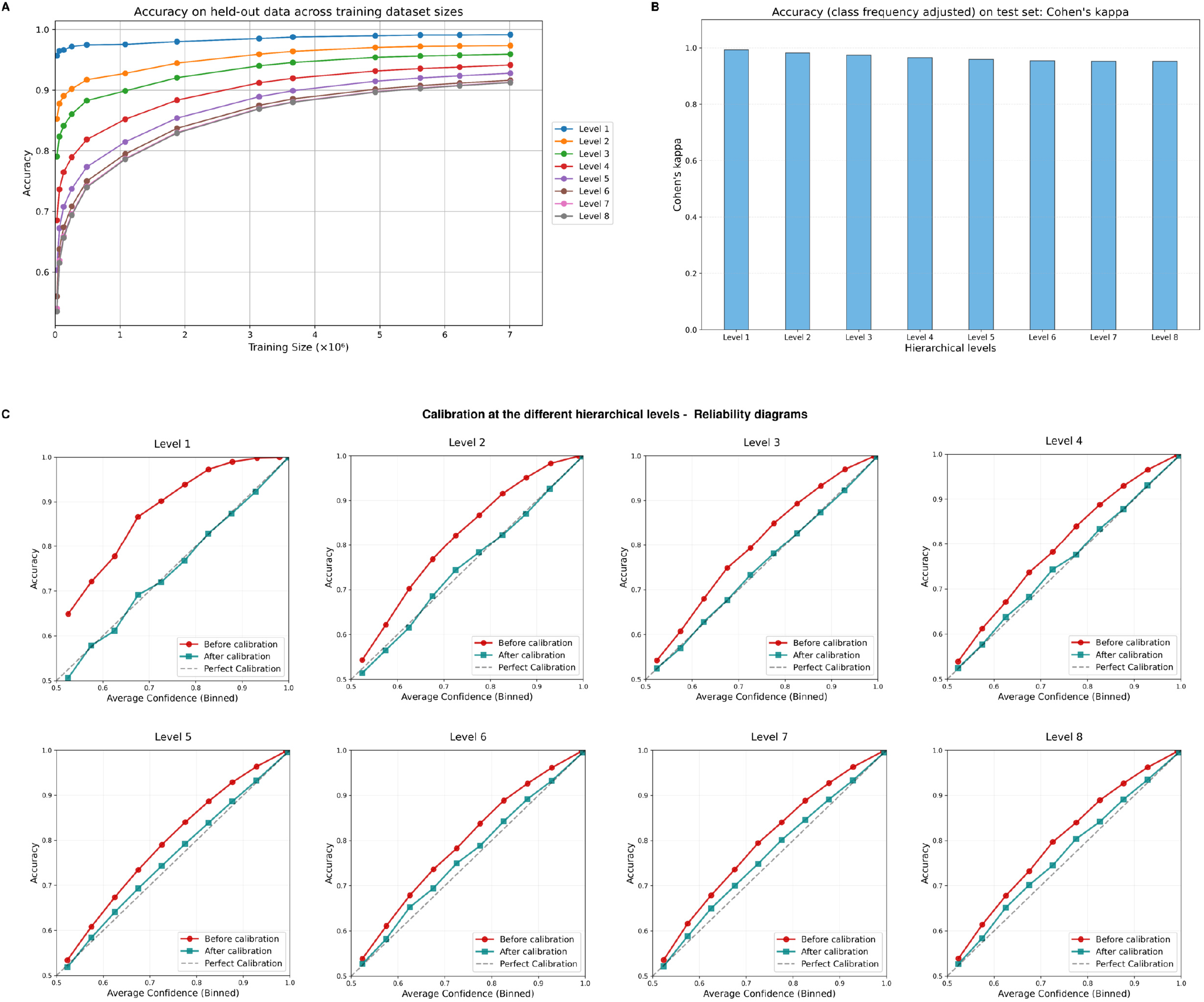
Training scale, class-balanced performance, and confidence calibration of Pan-human Azimuth. (A) Variation in validation accuracy of the Pan-human Azimuth architecture trained on datasets of increasing size. (B) Cohen’s kappa measured on the test set demonstrates that Pan-human Azimuth retains a high level of performance for the less abundant cell types and that model training is not saturated by the more abundant populations. (C) Accuracy (binned) as measured on the calibration test set before and after calibration plotted against average confidence in each bin at each hierarchical level. Measured accuracy and predicted confidence values are closely aligned post-calibration.

**Supplementary Figure 2.**
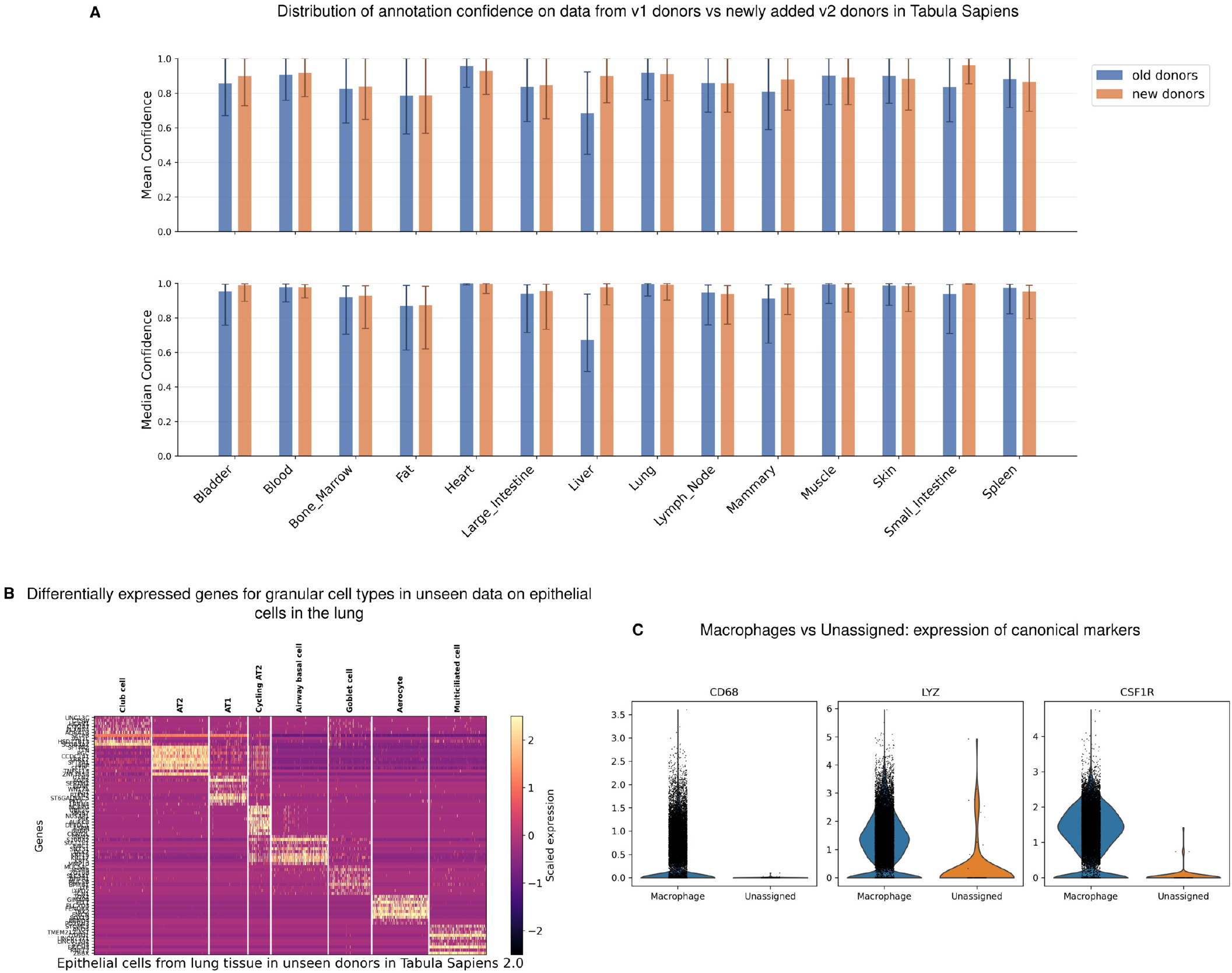
Additional analysis of model-derived annotations on Tabula Sapiens. (A) Pan-human Azimuth retains high annotation confidence on held-out data from new Tabula Sapiens v2 donors. Mean and median confidence (± standard deviation and interquartile range, respectively) across a range of tissues. (B) Fine grained annotations on epithelial cells from the lung are well supported by differential expression structure. (C) Accuracy of “Unassigned” calls by Pan-human Azimuth on cells manually annotated as macrophages: consensus macrophage calls show high expression of canonical markers *CD68, LYZ, CSF1R* while “Unassigned” cells generally do not.

**Supplementary Figure 3.**
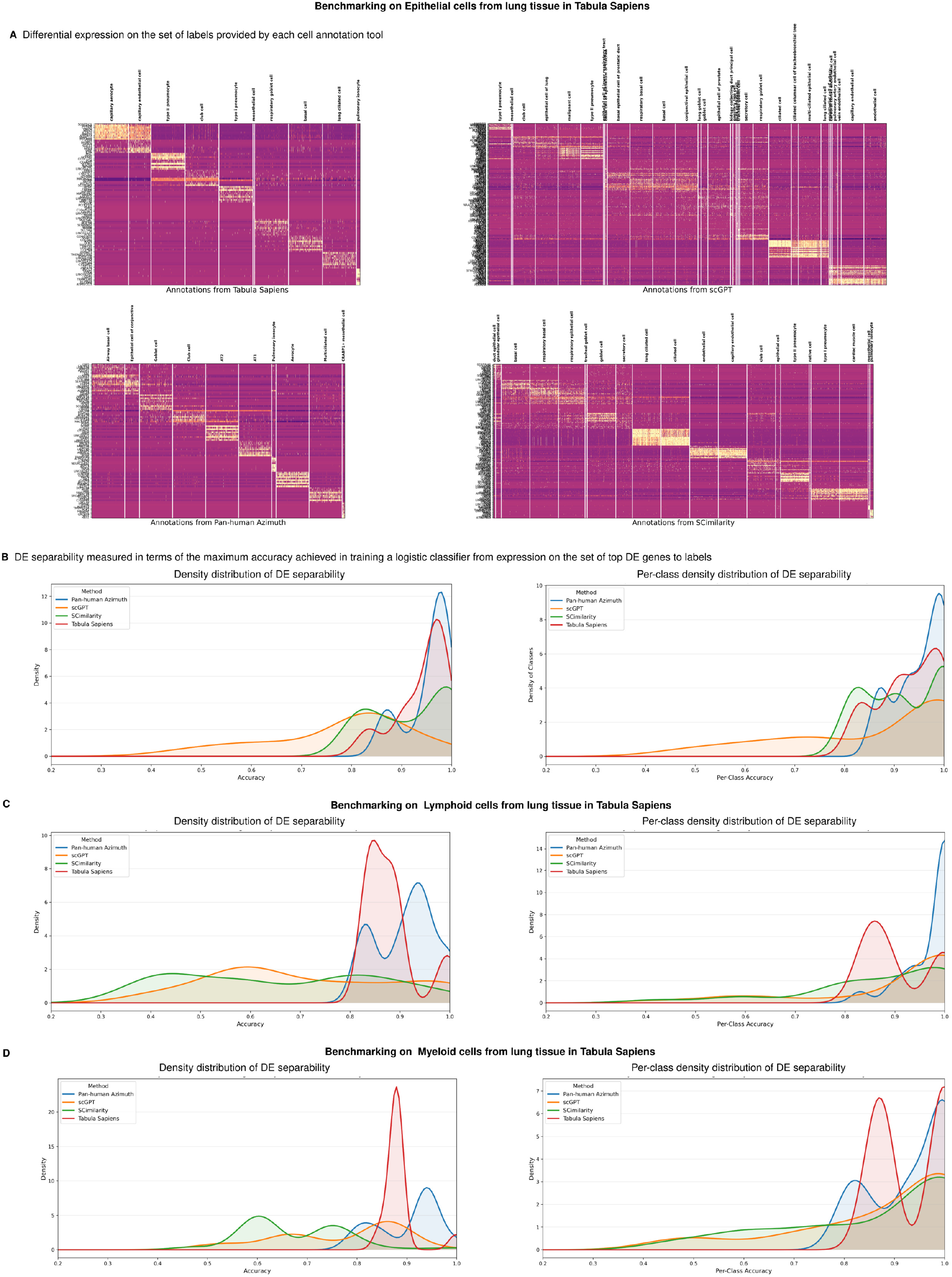
Benchmarking transcriptional separability of annotation labels. We quantify the transcriptional support of cell type annotations by assessing how well they are supported by differential gene expression using DE separability (Supplementary Methods). We compare Pan-human Azimuth against manual annotations from Tabula Sapiens, scGPT, and SCimilarity. (A) Differential-expression heatmaps for lung epithelial cells, grouped according to the labels assigned by each method: Tabula Sapiens manual annotations, scGPT, SCimilarity, and Pan-human Azimuth, shown clockwise from the top left. (B) Overall and per-class DE separability for lung epithelial cells. Pan-human Azimuth and the manual Tabula Sapiens annotations show similarly strong transcriptomic support, whereas labels assigned by scGPT and SCimilarity are less consistently separable, in agreement with the corresponding differential-expression heatmaps. (C) DE separability for lung lymphoid cells. Pan-human Azimuth labels show stronger transcriptomic support than those assigned by the other methods. (D) DE separability for lung myeloid cells, showing that Pan-human Azimuth labels are more consistently supported by distinguishable expression programs than labels from the other automated methods.

**Supplementary Figure 4.**
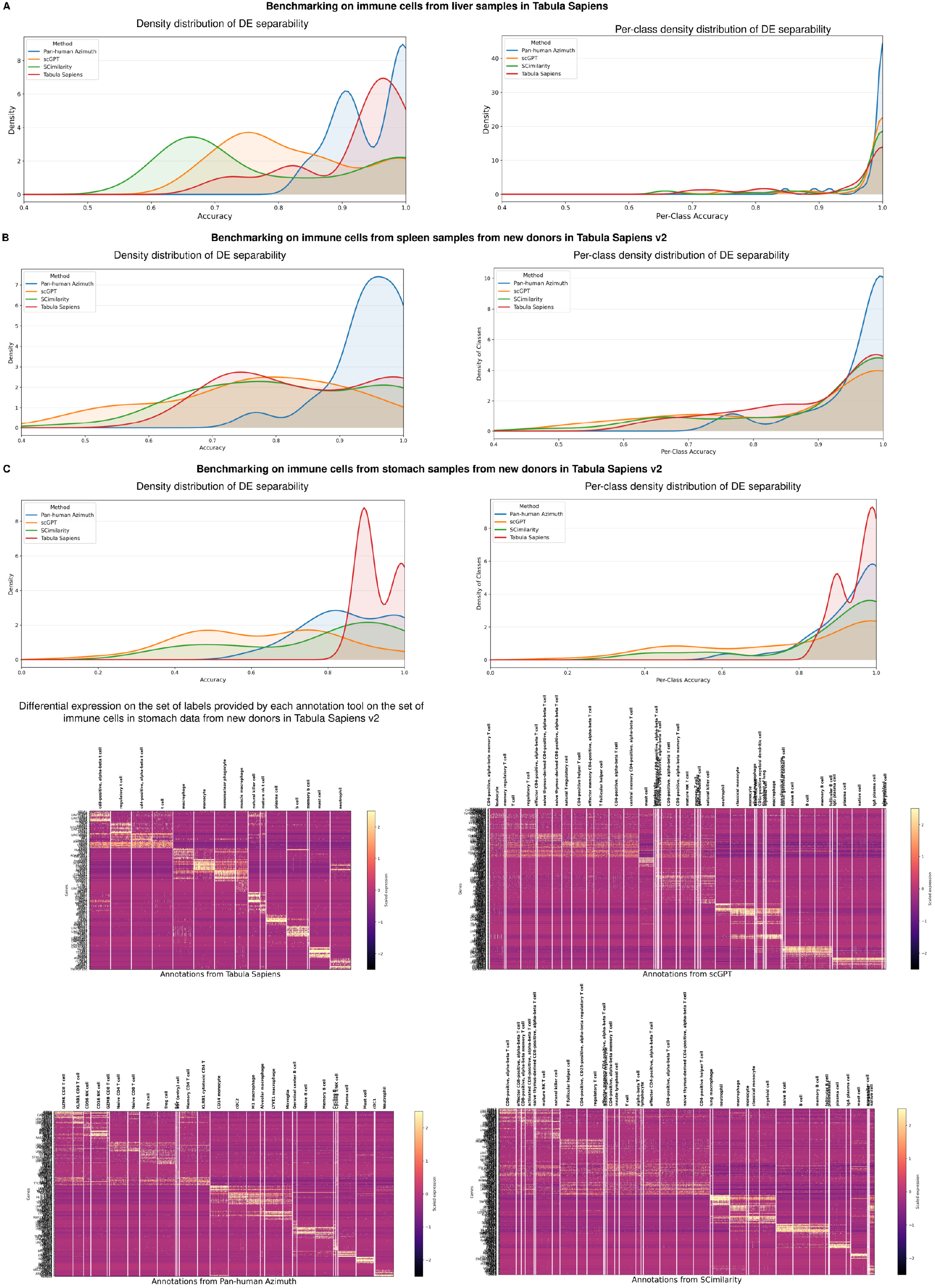
Quantifying transcriptional separability in additional tissues. (A) DE separability scores for immune cells from liver tissue in Tabula Sapiens v1. (B) DE separability quantification on immune cells from spleen tissue in Tabula Sapiens v2. Pan-human Azimuth outperforms all other methods in both overall and class-wise DE separability. (C) DE separability benchmarking on immune cells from stomach tissue in Tabula Sapiens v2. Tabula Sapiens manual annotations achieve the highest DE separability overall, while Pan-human Azimuth outperforms the other automated methods. DE heatmaps for stomach immune cells are consistent with the quantitative benchmarking results. Manual annotations show strong transcriptional support but annotate cells at lower resolution than automated methods, while annotations provided by scGPT and SCimilarity suffer from label fragmentation.

**Supplementary Figure 5.**
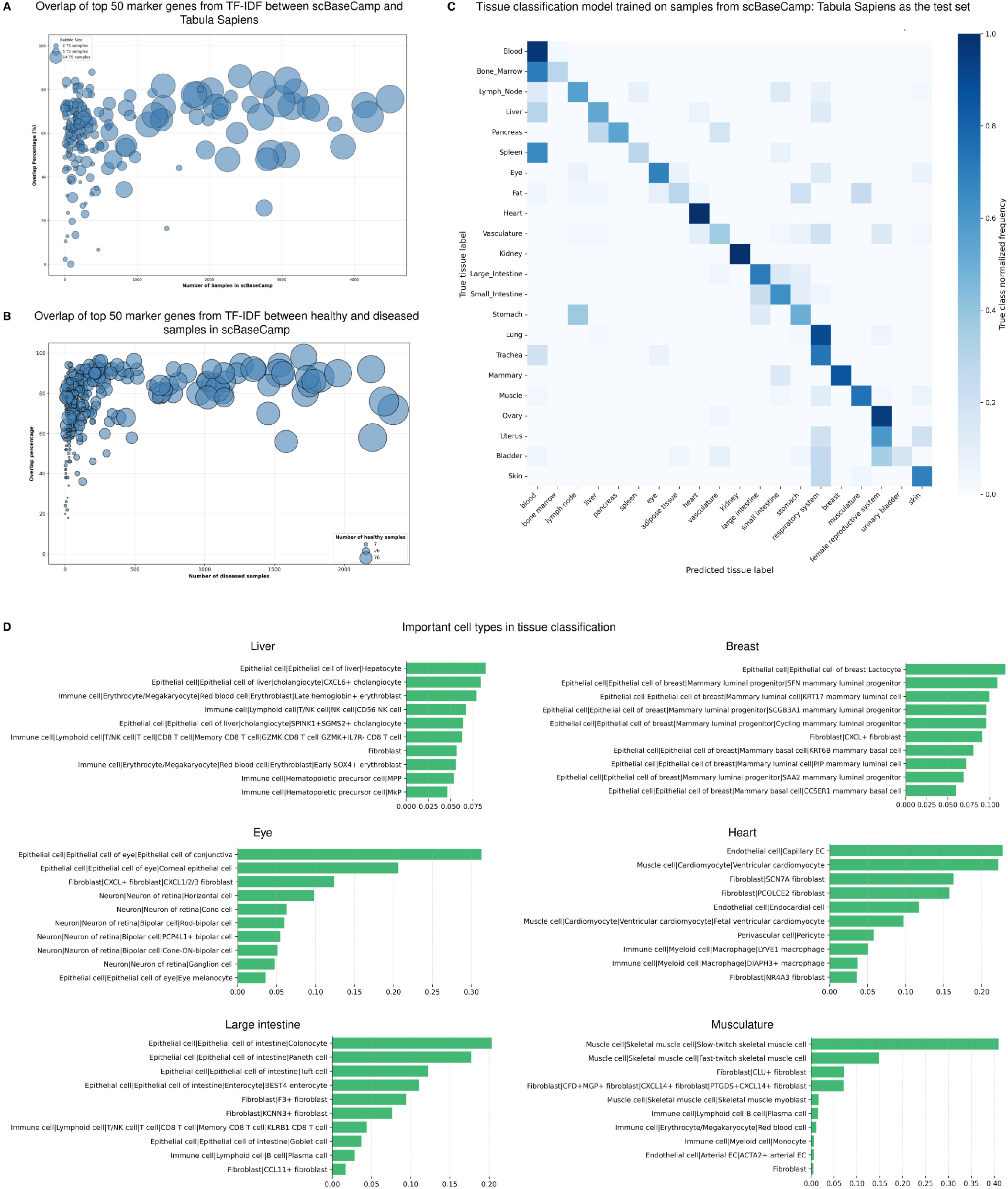
Cross-dataset marker reproducibility and tissue classification from cell type composition. (A) Overlap between the top 50 marker genes identified independently for shared Pan-human Azimuth cell types in scBaseCamp and Tabula Sapiens. Marker genes were derived using TF-IDF across non-cancer human samples, and the overlap supports the reproducibility of annotation-associated transcriptional programs across independent datasets. (B) Overlap between marker sets derived independently from healthy and diseased non-cancer scBaseCamp samples. The concordance of these marker programs indicates that many cell type-specific transcriptional signatures are preserved across disease contexts. (C) Confusion matrix evaluating a tissue classifier trained on scBaseCamp cell type proportions and applied to the independent Tabula Sapiens atlas. The strong diagonal structure shows that tissue-associated compositional patterns learned from scBaseCamp generalize across datasets. (D) Integrated Gradients attribution scores identifying the cell types that contribute most strongly to tissue classification. Epithelial populations are the dominant predictors for many tissues, while selected non-epithelial populations, including SCN7A^+^ fibroblasts in the heart, also exhibit strong tissue-associated contributions (see Fig.5E–F).

**Supplementary Figure 6.**
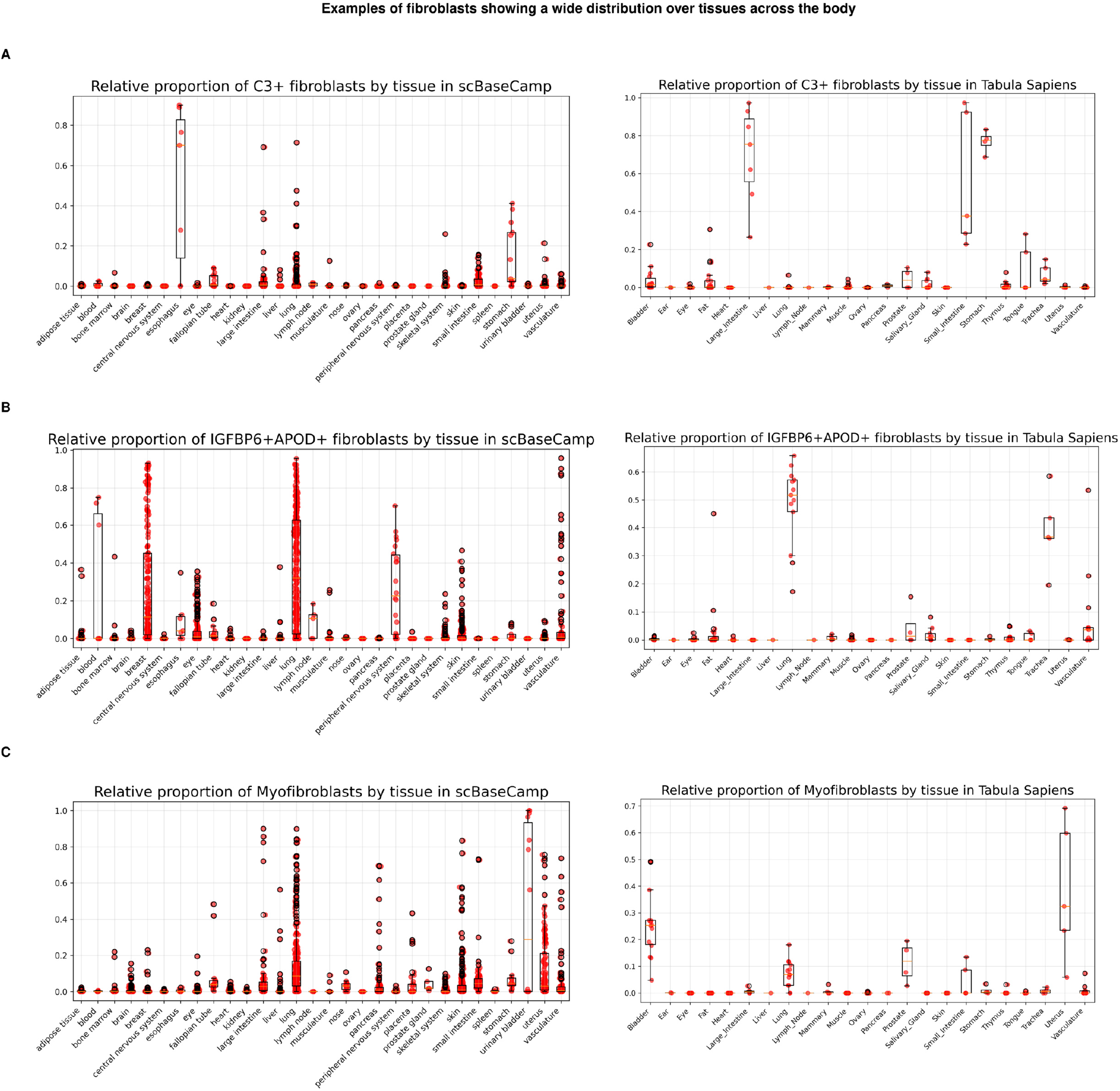
Fibroblast populations distributed across human tissues. Tissue distributions of (A) C3^+^ fibroblasts, (B) IGFBP6^+^APOD^+^ fibroblasts, and (C) myofibroblasts across scBaseCamp and Tabula Sapiens. Each population is detected across a broad range of tissues, and the overall distribution patterns are reproducible between the two independent datasets.

**Supplementary Figure 7.**
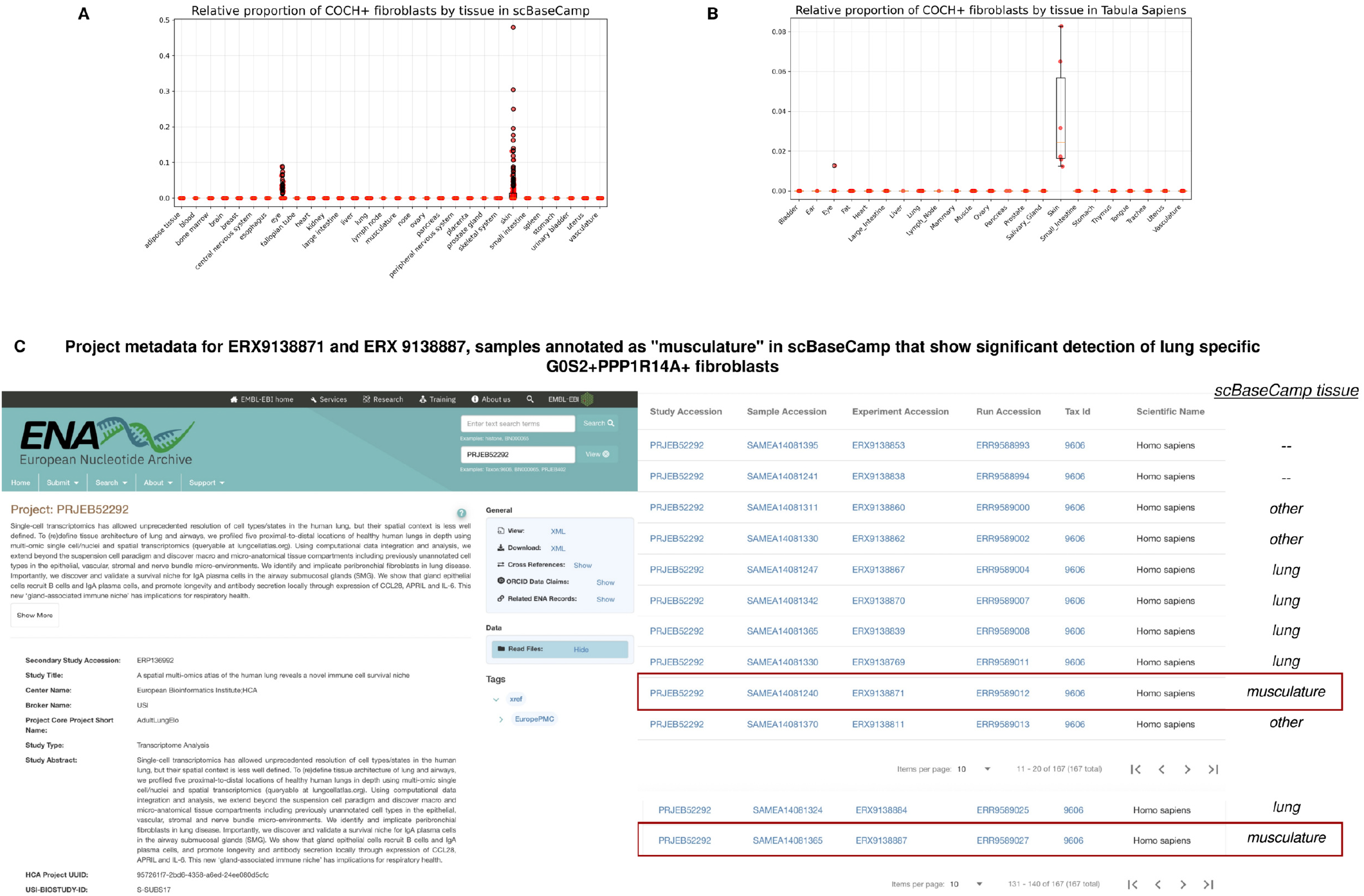
Tissue-restricted fibroblast populations and validation of tissue metadata. (A) Relative abundance of COCH^+^ fibroblasts across tissues in scBaseCamp, showing strong enrichment in skin and related ocular tissues, including the conjunctiva. (B) The skin-associated distribution of COCH^+^ fibroblasts is independently reproduced in Tabula Sapiens. (C) G0S2^+^PPP1R14A^+^ fibroblasts are strongly localized to the lung in both scBaseCamp and Tabula Sapiens, consistent with Fig.5C–D. The small number of apparent non-lung exceptions in scBaseCamp reflect errors in automated extraction of tissue metadata. For example, ERX9138871 and ERX9138887 were labeled as “musculature” in scBaseCamp but are identified as lung samples in the corresponding European Nucleotide Archive records for study PRJEB52292.

**Supplementary Figure 8.**
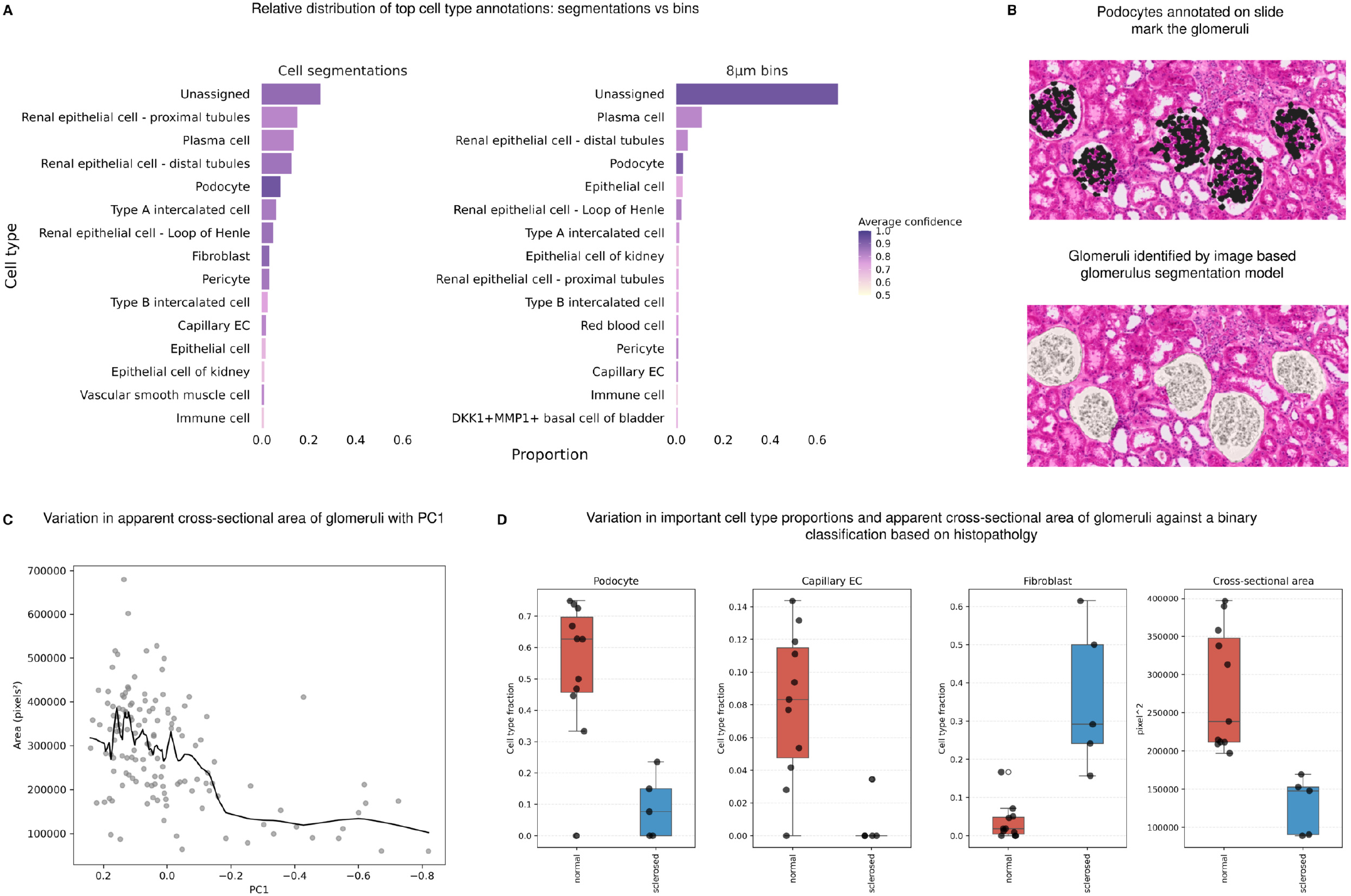
Segmentation-based annotation and glomerular heterogeneity in Visium HD kidney data. (A) Comparison of Pan-human Azimuth annotations obtained from segmentation-derived cell profiles and fixed 8 *µ*m bins. Segmentation-based profiles yield a substantially higher fraction of confidently assigned cells, consistent with improved recovery of single-cell molecular profiles relative to fixed-bin aggregation. (B) Spatial localization of Pan-human Azimuth-defined podocytes within the kidney cortex. Podocyte annotations coincide with visually apparent glomerular cross-sections and with glomerular masks generated independently by an image-based segmentation model trained on histopathology images. (C) Relationship between apparent glomerular cross-sectional area and PC1 derived from glomerular cell-type composition. (D) Comparison of podocyte, capillary endothelial-cell, and fibroblast-like fractions, apparent cross-sectional area, and PC1 between glomeruli classified by an expert renal pathologist as normal or sclerosed. The observed differences are consistent with the continuous variation in glomerular composition identified across the full set of segmented glomeruli.

**Supplementary Table 1.** Pan-human Azimuth neural network architecture. Layer-by-layer summary of the model architecture as returned by keras model.summary(). Each classification head (L1_out through L8_out) is preceded by a dense layer and dropout, and receives a concatenated input comprising the 128-dimensional embedding and compressed projections of all preceding levels’ softmax outputs.

| Layer | Type | Output dim | Parameters |
| --- | --- | --- | --- |
| *Embedding module* |  |  |  |
| input_layer | Input | 5,055 | 0 |
| dense | Dense | 1,024 | 5,177,344 |
| dense_1 | Dense | 512 | 524,800 |
| dense_2 | Dense | 256 | 131,328 |
| dense_3 | Dense | 128 | 32,896 |
| *Level 1 classification head (13 classes)* |  |  |  |
| dense_4 | Dense | 32 | 4,128 |
| dropout | Dropout | 32 | 0 |
| L1_out | Dense (softmax) | 13 | 429 |
| dense_5 | Dense (projection) | 3 | 42 |
| *Level 2 classification head (91 classes)* |  |  |  |
| concatenate | Concatenate | 131 | 0 |
| dense_6 | Dense | 64 | 8,448 |
| dropout_1 | Dropout | 64 | 0 |
| L2_out | Dense (softmax) | 91 | 5,915 |
| dense_7 | Dense (projection) | 22 | 2,024 |
| *Level 3 classification head (209 classes)* |  |  |  |
| concatenate_1 | Concatenate | 153 | 0 |
| dense_8 | Dense | 128 | 19,712 |
| dropout_2 | Dropout | 128 | 0 |
| L3_out | Dense (softmax) | 209 | 26,961 |
| dense_9 | Dense (projection) | 49 | 10,290 |
| *Level 4 classification head (313 classes)* |  |  |  |
| concatenate_2 | Concatenate | 202 | 0 |
| dense_10 | Dense | 256 | 51,968 |
| dropout_3 | Dropout | 256 | 0 |
| L4_out | Dense (softmax) | 313 | 80,441 |
| dense_11 | Dense (projection) | 71 | 22,294 |
| *Level 5 classification head (355 classes)* |  |  |  |
| concatenate_3 | Concatenate | 273 | 0 |
| dense_12 | Dense | 256 | 70,144 |
| dropout_4 | Dropout | 256 | 0 |
| L5_out | Dense (softmax) | 355 | 91,235 |
| dense_13 | Dense (projection) | 80 | 28,480 |
| *Level 6 classification head (374 classes)* |  |  |  |
| concatenate_4 | Concatenate | 353 | 0 |
| dense_14 | Dense | 256 | 90,624 |
| dropout_5 | Dropout | 256 | 0 |
| L6_out | Dense (softmax) | 374 | 96,118 |
| dense_15 | Dense (projection) | 84 | 31,500 |
| *Level 7 classification head (380 classes)* |  |  |  |
| concatenate_5 | Concatenate | 437 | 0 |
| dense_16 | Dense | 256 | 112,128 |
| dropout_6 | Dropout | 256 | 0 |
| L7_out | Dense (softmax) | 380 | 97,660 |
| dense_17 | Dense (projection) | 86 | 32,766 |
| *Level 8 classification head (382 classes)* |  |  |  |
| concatenate_6 | Concatenate | 523 | 0 |
| dense_18 | Dense | 256 | 134,144 |
| dropout_7 | Dropout | 256 | 0 |
| L8_out | Dense (softmax) | 382 | 98,174 |
| **Total** |  |  | **6,981,993** |

